# Systematic assessment of tissue dissociation and storage biases in single-cell and single-nucleus RNA-seq workflows

**DOI:** 10.1101/832444

**Authors:** Elena Denisenko, Belinda B. Guo, Matthew Jones, Rui Hou, Leanne de Kock, Timo Lassmann, Daniel Poppe, Olivier Clement, Rebecca K. Simmons, Ryan Lister, Alistair R. R. Forrest

## Abstract

Single-cell and single-nucleus RNA sequencing have been widely adopted in studies of heterogeneous tissues to estimate their cellular composition and obtain transcriptional profiles of individual cells. However, the current fragmentary understanding of artefacts introduced by sample preparation protocols impedes the selection of optimal workflows and compromises data interpretation. To bridge this gap, we compared performance of several workflows applied to adult mouse kidneys. Our study encompasses two tissue dissociation protocols, two cell preservation methods, bulk tissue RNA sequencing, single-cell and three single-nucleus RNA sequencing workflows for the 10x Genomics Chromium platform. These experiments enable a systematic comparison of recovered cell types and their transcriptional profiles across the workflows and highlight protocol-specific biases important for the experimental design and data interpretation.

## Introduction

Single-cell RNA sequencing (scRNA-seq) is an increasingly powerful technology that enables analysis of gene expression in individual cells. ScRNA-seq has been recently used to study organism development [1–3], normal tissues [4–6], cancer [7–10] and other diseases [11, 12]. These studies have shed light on tissue heterogeneity and provided previously inaccessible insights into tissue functioning.

Advances in high-throughput droplet-based microfluidics technologies have facilitated analysis of thousands of cells in parallel [13–15], and Chromium from 10x Genomics has become a widely used commercial platform [15]. Multiple tissue preparation protocols are compatible with Chromium, but the protocol of choice should ideally maintain RNA integrity and cell composition of the original tissue.

Solid tissues need to be dissociated to release individual cells suitable for 10x Genomics Chromium scRNA-seq. However, optimal dissociation needs to achieve a balance between releasing cell types that are difficult to dissociate while avoiding damage to those that are fragile. Tissue dissociation is most commonly conducted using enzymes which require incubation at 37°C for variable times based on tissue type. At this temperature, the cell transcriptional machinery is active, hence, gene expression can be altered in response to the dissociation and other environmental stresses [16, 17]. A recent alternative approach minimising this artefact uses cold-active protease to conduct tissue dissociation on ice [18]. Alternatively, single-nucleus RNA sequencing protocols (snRNA-seq) use much harsher conditions to release nuclei from tissue and can be applied to snap frozen samples, thus avoiding many of the dissociation-related artefacts [19, 20]. Single nuclei methods should also permit profiling of nuclei from large cells (>40 µm) that do not fit through the microfluidics.

Additional restrictions and challenges are faced by complex experimental designs where specimens cannot be processed immediately. In this case, samples need to be preserved either as an intact tissue or in a dissociated form as a single-cell suspension. Each of the approaches mentioned above introduces specific biases and artefacts that can manifest themselves in altered transcriptional profiles or altered representation of cell types. These biases need to be considered when designing and analysing data from a single-cell experiment; however, they are still incompletely understood.

Some of the artefacts have been investigated in recent studies comparing single-cell profiles of methanol-fixed and live cells [21, 22], cryopreserved and live cells [22, 23], single-cell and single-nucleus protocols [24–26], or tissue dissociation using cold-active protease and traditional digestion at 37°C [18]. However, these assessments were performed in different tissues under different conditions and lack extensive comparison to bulk data.

Here, we performed a comprehensive study in healthy adult mouse kidneys using 10x Genomics Chromium workflows for scRNA-seq and snRNA-seq, along with bulk RNA-seq of undissociated and dissociated tissue. We compare and contrast two tissue dissociation protocols (digestion at 37°C, further referred to as warm dissociation, or with cold-active protease, further referred to as cold dissociation), two single-cell suspension preservation methods (methanol fixation and cryopreservation) and three single nuclei isolation protocols (**Fig. 1**). A total of 77,656 single-cell, 98,303 single-nucleus and 15 bulk RNA-seq profiles were generated and made publicly available (GSE141115). Our dissection of artefacts associated with each of the approaches will serve as a valuable resource to aid interpretation of single-cell and single-nucleus gene expression data and help to guide the choice of experimental workflows.

**Figure 1.**
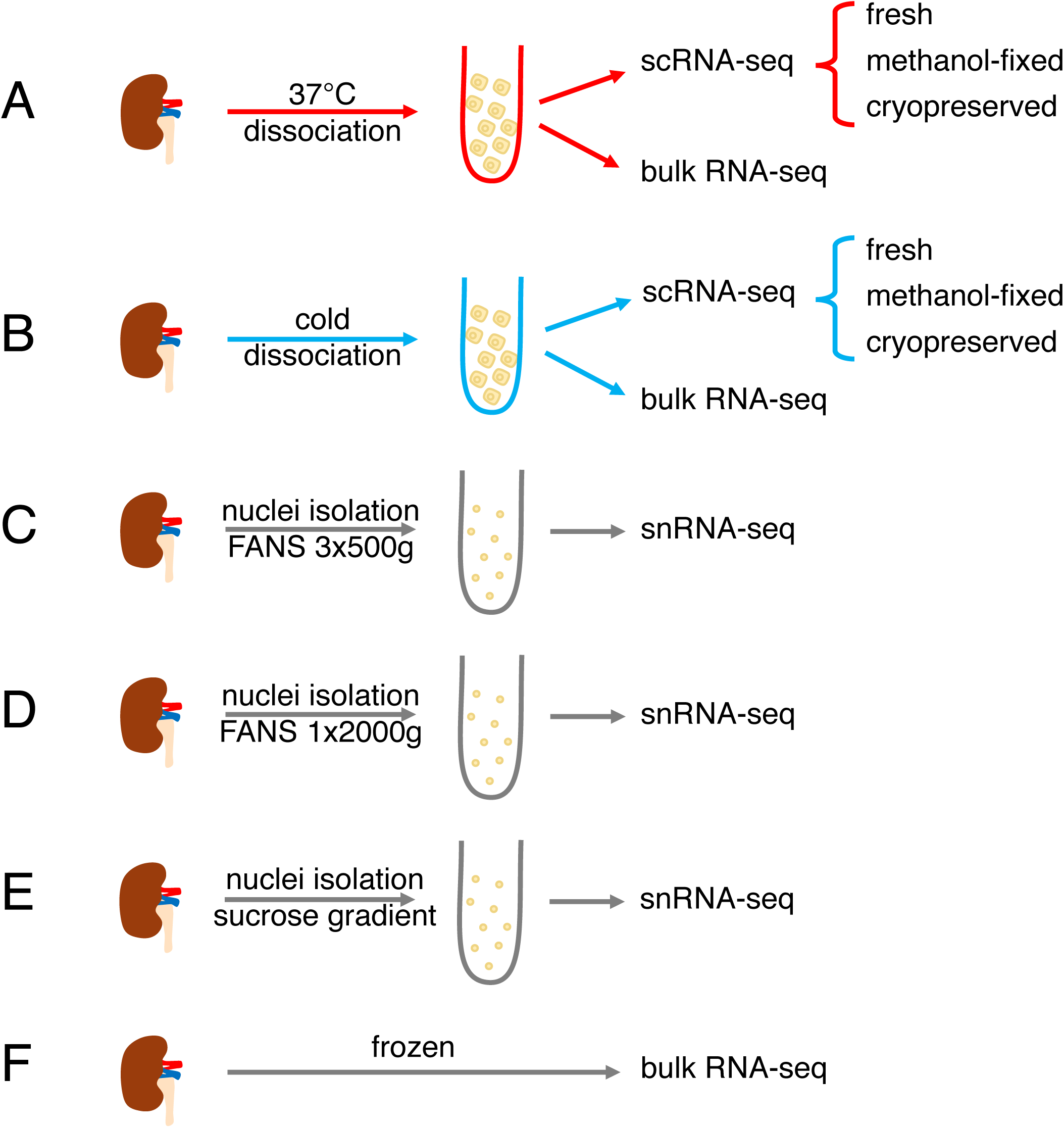
Overview of experiments performed in this study. All experiments were carried out in biological triplicate using three kidneys from three different mice. **a**) 37°C dissociation used the Multi-tissue dissociation kit 2 from Miltenyi Biotec. **b**) Cold dissociation was carried out on ice using *B. Licheniformis* protease. In **a** and **b**, methanol fixed samples used 80% MeOH at -20°C and then were stored at -80°C. Cryopreservation was carried out using 50% FBS, 40% RPMI-1640, 10% DMSO with gradient cooling to -80°C then storage in liquid nitrogen. **c-e**) Whole kidneys were flash frozen using an isopentane bath -30°C and then stored at -80°C. Three different nuclei preparation methods were tested using either fluorescently activated nuclei sorting (FANS) or a sucrose gradient to enrich for singlet nuclei. **f**) Bulk RNA-seq was carried out using the NEBNext Ultra II RNA Library Kit for Illumina with rRNA depletion or NEBNext Poly(A) mRNA isolation module. See **Methods** and **Supplementary Table 17** for more details.

## Results

### Comparison of tissue dissociation protocols

In the first series of experiments, we set out to compare two tissue dissociation protocols using kidneys from adult male C57BL/6J mice. Kidneys were dissociated at 37°C using a commercial Miltenyi Multi Tissue Dissociation Kit 2 or on ice using a cold-active protease from *Bacillus licheniformis* (**Methods**, **Fig. 1A, B**). Aliquots of single cell suspensions were profiled using 10x Genomics Chromium scRNA-seq and a bulk RNA-seq protocol (**Methods**, **Fig. 1A, B**). All experiments were performed in triplicate and data were processed as described in **Methods**.

#### Warm tissue dissociation induces stress response

Bulk RNA-seq profiling of single-cell suspensions revealed induction of stress response genes in warm-dissociated samples. Differential expression analysis identified 71 genes with higher expression in warm-dissociated kidneys and 5 genes with higher expression in cold-dissociated kidneys (logFC > 2, FDR < 0.05, edgeR exact test [27], **Supplementary Table 1**). Gene ontology analysis with ToppGene [28] reported “regulation of cell death” as the top significantly enriched Biological Process for the genes more highly expressed in warm-dissociated kidneys (an overlap of 22 genes, FDR = 1.7E-7, see **Supplementary Table 1**). Genes with the highest logFC values (>4) included immediate-early genes *Fosb, Fos, Jun, Junb, Atf3, Egr1* and heat shock proteins *Hspa1a* and *Hspa1b* (**Fig. 2A**). These findings from bulk RNA-seq confirm the original observations of Adam *et al.* [18] that warm tissue dissociation induces substantial stress-response-related changes.

**Figure 2.**
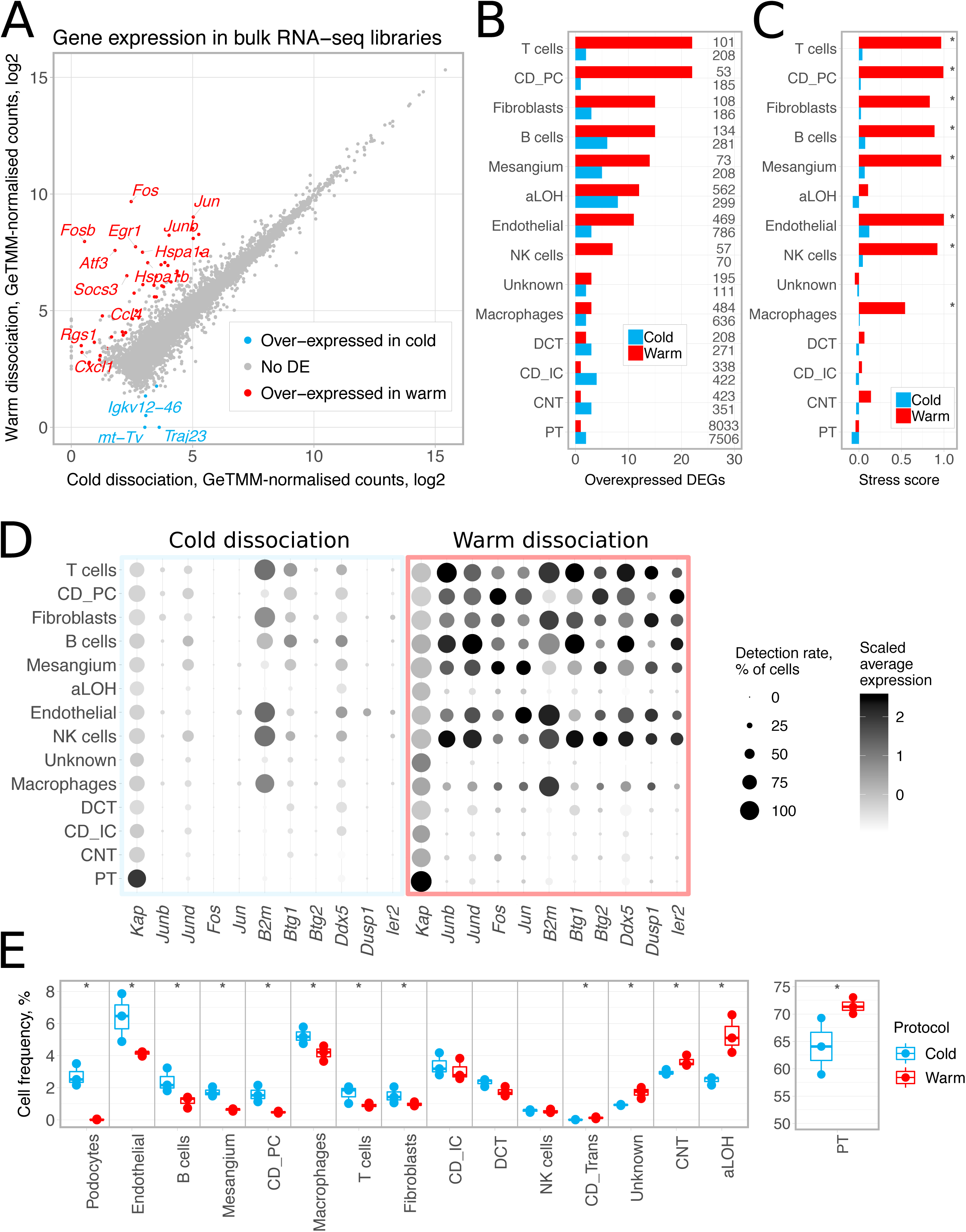
Comparison of cold and warm tissue dissociation protocols. **a**) Bulk RNA-seq profiles of dissociated kidneys. GeTMM-normalised counts [43] were averaged across three biological replicates and log2-transformed after adding a pseudo count of 1. DEGs with FDR < 0.05 and logFC threshold of 2 (edgeR exact test [27]) are shown as red and blue dots; protein-coding genes with logFC > 4 are labelled. **b**) Number of differentially expressed genes (DEGs) between cold- and warm-dissociated scRNA-seq libraries. Calculated for each cell type separately using Wilcoxon test in Seurat [34] with thresholds of logFC = 0.5, minimum detection rate 0.5, FDR < 0.05. Numbers on the right side of the plot indicate cell population size. **c**) Stress score – an expression score for a set of 17 stress-response-related genes (*Fosb, Fos, Jun, Junb, Jund, Atf3, Egr1, Hspa1a, Hspa1b, Hsp90ab1, Hspa8, Hspb1, Ier3, Ier2, Btg1, Btg2, Dusp1*). Calculated as average gene expression level of these genes subtracted by averaged expression of randomly selected control genes and then averaged for cell types. Significance was calculated in a Monte-Carlo procedure with 1000 randomly selected gene sets of the same size, asterisks denote p-value < 0.01. **d**) Expression and detection rates of differentially expressed genes commonly induced in warm-dissociated samples (differentially expressed in at least four cell types). **e**) Cell type composition of freshly profiled scRNA-seq libraries. Three biological replicates are shown per condition. Asterisks denote two-sided chi-square test p-value < 0.001. In **b-d**, podocytes and transitional cells were excluded due to low cell numbers. aLOH: ascending loop of Henle; CD_IC: intercalated cells of collecting duct; CD_PC: principal cells of collecting duct; CNT: connecting tubule; DCT: distal convoluted tubule; PT: proximal tubule.

#### Single-cell sequencing reveals heterogeneous stress response across cell populations

We next characterised differences between the two tissue dissociation protocols by scRNA-seq profiling of fresh cell suspensions (**Fig. 1A, B**). This dataset comprised 23,108 cells, including 11,851 cells from cold- and 11,257 cells from warm-dissociated kidneys (**Methods**). Cells were classified into 15 cell types using scMatch [29] by comparing their expression to reference expression profiles from three previous mouse kidney studies [26, 30, 31], followed by gene signature-based refinement (**Methods**, **Supplementary Fig. 1-3**, **Supplementary Table 2**).

Differential expression analysis identified 64 genes more highly expressed in warm-dissociated libraries in at least one cell type (**Fig. 2B**, **Supplementary Table 3**) and gene ontology analysis again reported “regulation of cell death” as one of the top significantly enriched terms (an overlap of 23 genes, FDR = 3.9E-7, **Supplementary Table 3**). The genes most commonly over-expressed across cell populations are shown in **Fig. 2D** and include immediate-early response genes such as *Junb* and *Jund* (differentially expressed in seven cell types) and *Jun* and *Fos* (differentially expressed in five cell types).

Notably, the numbers of differentially expressed genes varied among cell types (**Fig. 2B**), suggesting that cell types respond differently to warm tissue dissociation. To quantify these differences, we selected a set of 17 known stress-response-related genes that were induced in the warm-dissociated samples (*Fosb, Fos, Jun, Junb, Jund, Atf3, Egr1, Hspa1a, Hspa1b, Hsp90ab1, Hspa8, Hspb1, Ier3, Ier2, Btg1, Btg2, Dusp1*) and used them to calculate a stress score (see **Methods**). **Fig. 2C** shows that significantly high stress scores were detected only in warm-dissociated samples in eight out of 14 cell types. Taken together, these results highlight that certain cell types, such as immune and endothelial cells, are particularly sensitive to warm tissue dissociation.

In contrast to the 64 genes with higher expression in the warm dissociation, only 20 genes had higher expression in the cold-dissociated cell populations, and only five of them (*Hbb-bs, Hba-a1, Hba-a2, mt-Co1, Malat1*) were identified in at least two cell types (**Fig. 2B**, **Supplementary Table 4**). We note the levels of haemoglobin transcripts suggest contamination from erythrocytes is higher in the samples dissociated on ice.

#### Cell composition differs between two tissue dissociation protocols

In addition to expression changes, our analyses identified eight cell populations that were less abundant in warm-dissociated samples in comparison to cold-dissociated ones, including podocytes, mesangial cells, and endothelial cells (**Fig. 2E left**, chi-square test p-value < 0.001). These depleted populations also showed significantly high expression of the stress-response-related gene set as described above (**Fig. 2C**). Notably, only three podocytes were detected in warm-dissociated samples (0.03% of the total cell count), compared to 330 (2.78%) in the cold-dissociated samples. These findings suggest that these populations are sensitive to warm dissociation and consequently underrepresented.

Conversely, we identified cells such as those of the ascending loop of Henle (aLOH) and proximal tubule (PT), that were more abundant in warm-dissociated samples (aLOH 4.99% vs. 2.52% in cold, PT 71.36% vs. 63.34% in cold), potentially indicating their less efficient dissociation by cold-active protease (**Fig. 2E right**). The bulk RNA-seq data suggest that similar proportions of aLOH cells still make it into suspension and that they might in fact be lost in the microfluidic partitioning (**Supplementary Note 1**).

### Comparison of cell preservation protocols

We next evaluated whether cryopreservation and methanol fixation maintain cell composition and transcriptional profiles of kidneys. Aliquots of single-cell suspensions of cold- and warm-dissociated kidneys were cryopreserved (50% FBS, 40% RPMI-1640, 10% DMSO) and stored for 6 weeks or methanol-fixed and stored for 3 months. These stored samples were then profiled with 10x Genomics Chromium scRNA-seq (**Fig. 1A, B**, **Methods**). The resulting datasets consisted of 11,627 and 5,545 methanol-fixed cells and 3,519 and 3,483 cryopreserved cells derived from cold- and warm-dissociated kidneys, respectively. Notably, only ∼32% of cells were recovered after cryostorage and the average viability estimated by the Countess was 75%. Despite loading similar numbers of cells, the number of high-quality cells obtained from the cryopreserved samples after quality control and filtering (**Methods**) was substantially lower (∼30%) than that of the fresh and methanol fixed samples.

#### Cryopreservation depletes epithelial cell types

The most prominent difference in recovery rates pertained to cells of the proximal tubule (PT), the most populous cell type in kidney [32]. In freshly profiled suspensions, PT composed 63.12% and 70.86% of all cells in cold- and warm-dissociated samples, respectively. In contrast, PT were scarcely detected in cryopreserved samples, at 0.31% and 0.57%, respectively (see **Fig. 3A** for cold-dissociated samples, **Supplementary Fig. 5** for warm-dissociated samples, **Supplementary Fig. 6** for biological replicates). We next compared recovery rates of other cell populations in freshly profiled and cryopreserved samples relative to all non-PT cells. This comparison revealed significant underrepresentation (chi-square test p-value < 0.001) of five kidney cell types in cryopreserved samples prepared with the cold dissociation protocol, three of which were also underrepresented in the cryopreserved warm dissociation samples (**Supplementary Fig. 7**). Together with the loss of PT cells, this indicates that the cryopreservation and subsequent thawing protocol failed to efficiently recover kidney epithelial cell populations.

**Figure 3.**
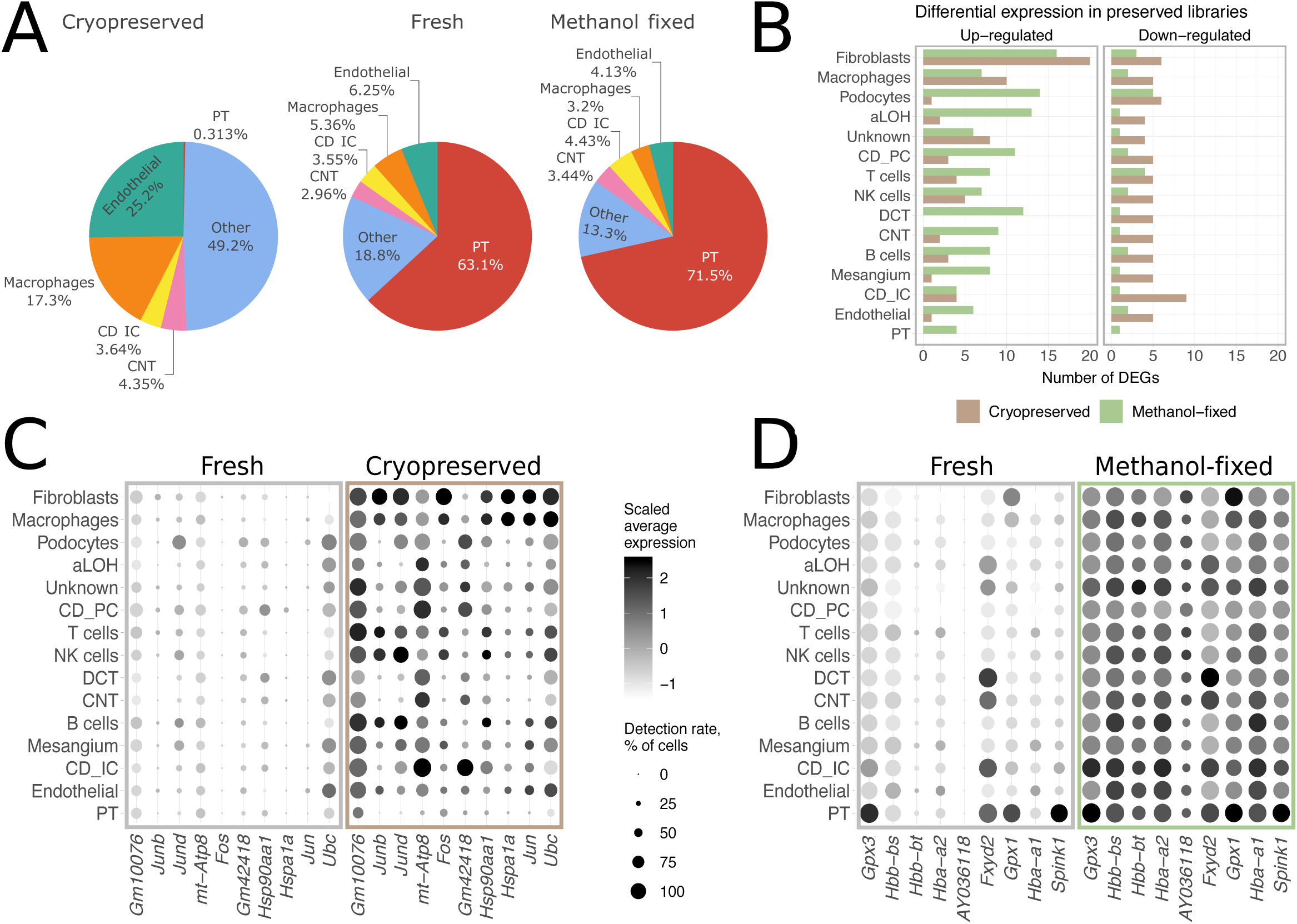
Cell preservation protocol performance in cold-dissociated samples. **a**) Cell type composition of freshly profiled and preserved cold-dissociated samples. **b**) Number of differentially expressed genes (DEGs) detected between preserved and freshly profiled aliquots. Seurat Wilcoxon test [34] with logFC = 1, min detection rate 0.5, FDR < 0.05 as thresholds. **c**) Expression and detection rates of DEGs with higher expression in cryopreserved samples in at least two cell types. **d**) Expression and detection rates of DEGs with higher expression in methanol-fixed samples in at least nine cell types. aLOH: ascending loop of Henle; CD_IC: intercalated cells of collecting duct; CD_PC: principal cells of collecting duct; CNT: connecting tubule; DCT: distal convoluted tubule; PT: proximal tubule.

Previous studies have reported cryopreserved cells generate comparable data to that of fresh cells [22, 23]. Hence, we repeated the experiment comparing cryopreserved and freshly profiled cold-dissociated single-cell suspension aliquots using different mice (Balb/c female), 10x chemistry (v3 as opposed to v2), storage length (2 weeks as opposed to 6 months) and centrifugation speed for thawing and resuspension (1200g as opposed to 400g). Again, there was a significant depletion of PT cells, with them making up 55.55% of the freshly profiled cells but only 7.65% of the cryopreserved cells (**Supplementary Fig. 8**). From this, we conclude that, at least in the case of mouse kidneys, cryopreservation of dissociated cells using 50% FBS, 40% RPMI, 10% DMSO can induce substantial deleterious changes in cell composition.

In contrast to a previous report assessing storage of cell lines and immune cells [22], in the case of dissociated mouse kidneys, methanol fixation better preserved cell type composition than cryopreservation (**Fig. 3A**, **Supplementary Fig. 5**, **Supplementary Fig. 6**). Nevertheless, certain cell types were moderately under-represented in the methanol-fixed samples in comparison to freshly profiled samples, with macrophages showing the largest reduction from 5.36% to 3.2% in cold-dissociated samples and from 4.28% to 2.54% in warm-dissociated samples.

#### Cryopreservation induces stress response

To gain further insights into preservation-related artefacts, we compared gene expression between preserved and freshly profiled samples in each cell type separately. In cold-dissociated samples, 31 and 27 genes were over-expressed in at least one cell type in cryopreserved and methanol-fixed cells, respectively, when compared to freshly profiled suspensions (**Fig. 3B**, see **Supplementary Fig. 5** for warm-dissociated samples).

In cryopreserved samples, stress response-related genes were induced, including multiple immediate-early response genes and heat shock proteins (**Fig. 3C**, **Supplementary Fig. 5**, **Supplementary Table 5-6**). In contrast, genes over-expressed in methanol-fixed cells were those highly expressed in tubular cells and haemoglobin genes (**Fig. 3D**, **Supplementary Table 7-8**). The same set of transcripts contaminated most cell types suggesting methanol fixation damages cells and leads to ambient RNA contamination of droplets.

### Comparison of single-cell and single-nucleus sequencing protocols

Having identified cold-active protease as a less damaging tissue dissociation approach for scRNA-seq, we next compared it to snRNA-seq. We performed a series of experiments using kidneys from Balb/c male mice with v2 10x chemistry or female mice with v3 chemistry and prepared cells using cold tissue dissociation for scRNA-seq and nuclei using three variant protocols for snRNA-seq (**Methods, Fig. 1B-E**). Two nuclei isolation protocols made use of fluorescence activated nuclei sorting (FANS). The first protocol washed the nuclei three times and used a centrifugation speed of 500g (further referred to as SN_FANS_3x500g, **Fig. 1C**). In the second protocol, nuclei were washed once and a centrifugation speed of 2000g was used (SN_FANS_1x2000g, **Fig. 1D**). In the third protocol, nuclei were initially washed using a 500g spin and then cleaned using a sucrose cushion avoiding the requirement to sort isolated nuclei (SN_sucrose, **Fig. 1E**). The three nuclei isolation protocols yielded comparable results, with the most notable difference being a higher contamination with mitochondrial genes in SN_FANS_1x2000g (**Supplementary Note 2**). In addition, we performed bulk RNA-seq of intact flash-frozen whole kidneys and of cold-dissociated cell suspensions (**Fig. 1B, F**).

Detection rates of non-epithelial kidney cell types were markedly different between scRNA-seq and snRNA-seq libraries (**Fig. 4A**, **Supplementary Fig. 9**, **Supplementary Table 9**). Immune cells were detected at lower rates in snRNA-seq (average of 0.73%) than in scRNA-seq (average of 6.03%) across all experiments performed (**Fig. 4A**, **Supplementary Fig. 9**).

**Figure 4.**
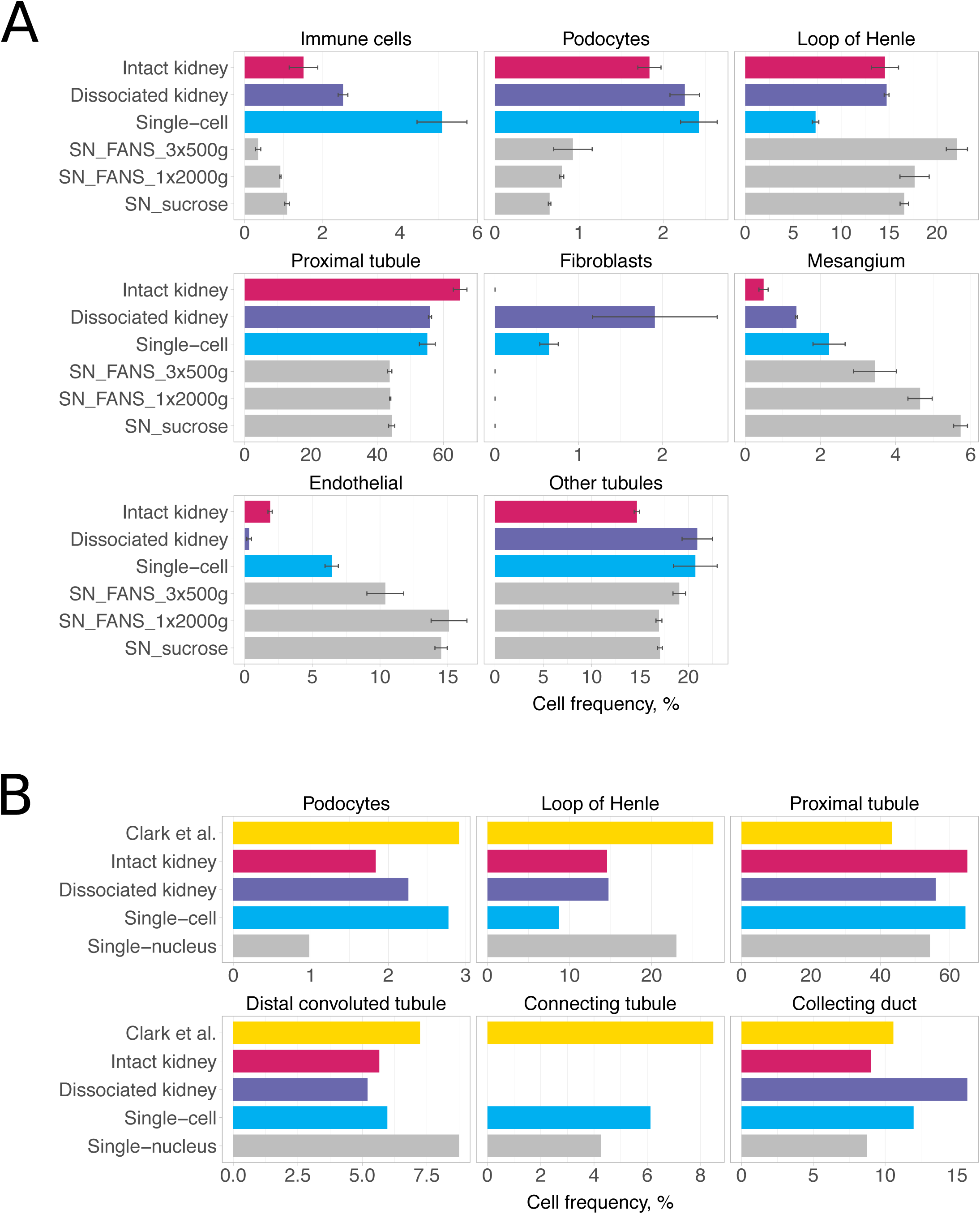
Comparison of single-cell and single-nucleus libraries. **a**) Cell type composition for kidneys from Balb/c female mice. Average percentages for scRNA-seq libraries are shown in blue and for snRNA-seq libraries in grey. BSEQ-sc estimates are shown for bulk RNA-seq of intact and dissociated kidneys. Error bars are standard error of mean. **b**) Abundance of renal epithelial cell types in Clark *et al*. study [32] in comparison to our data from Balb/c female mice.

Using the bulk RNA-seq from intact kidneys (**Fig. 1F**) and BSEQ-sc [33] to predict the proportions of each cell type present, we estimated that approximately 1.51% should correspond to immune cells in Balb/c female mice and 4.84% in Balb/c male mice. This suggests an underrepresentation of immune cells in the snRNA-seq data. Furthermore, macrophages were the only type of immune cells recovered in snRNA-seq libraries, whereas in scRNA-seq libraries we also detected T cells (1.38% on average), B cells (0.77%) and NK cells (0.65%). Similarly, podocytes composed only 0.7% in snRNA-seq libraries as opposed to 3.28% in scRNA-seq (**Fig. 4A**, **Supplementary Fig. 9**). Cell types more abundant in snRNA-seq libraries included loop of Henle and mesangial cells (**Fig. 4A**, **Supplementary Fig. 9**).

We next compared the observed cell composition to estimates of epithelial cell type contribution based on quantitative renal anatomy, as reported by Clark *et al.* recently [32]. **Fig. 4B** and **Supplementary Fig. 9** show that for some cell types, such as podocytes, scRNA-seq yields proportions most similar to the quantitative renal anatomy estimates, whereas for other cell types, such as loop of Henle cells, snRNA-seq better captures cell composition. Bulk RNA-seq-based proportions estimated from intact kidneys largely contradicted the anatomical estimates, which might reflect inaccurate deconvolution of the sample. Finally, comparison of bulk RNA-seq profiles of intact kidneys and cold-dissociated cell suspensions from female Balb/c mice suggested cell types that may be unequally represented in whole vs dissociated kidneys (**Supplementary Note 3**).

Differential expression analysis comparing individual cell types profiled by snRNA-seq and scRNA-seq suggested higher expression of long noncoding RNAs in snRNA-seq libraries and higher expression of genes related to mitochondrial and ribosomal functions in scRNA-seq, in agreement with previous reports [24, 26] (**Supplementary Table 10**).

Finally, as mitotic cells lack a nuclear membrane and in principle should not be observed in the snRNA-seq data, we inferred cell cycle phases for cells and nuclei using Seurat (**Methods**) [34]. Notably, Seurat predicted a higher fraction of G1 phase cells and lower fraction of S phase cells in scRNA-seq libraries when compared to snRNA-seq libraries for virtually all cell types (**Supplementary Fig. 10Supplementary Fig. 9**). This suggests that there are indeed underlying biases in cell cycle phase distributions in snRNA-seq data in comparison to scRNA-seq data; however, to fully dissect this, a classifier that can discriminate mitotic cells from early G1 and late G2 is required.

## Discussion

Interrogating complex tissues at the level of individual cells is essential to understand organ development, homeostasis and pathological changes. Despite the rapid advancement and widespread adoption of scRNA-seq and snRNA-seq technologies, the associated biases remain incompletely understood. To characterise some of the biases, we performed a systematic comparison of recovered cell types and their transcriptional profiles across two tissue dissociation protocols, two single-cell suspension preservation methods and three single-nuclei isolation protocols, each with three biological replicates per experiment.

Previous studies have reported on artefactual gene expression changes induced by proteolytic tissue digestion at 37°C in sensitive cell populations [16, 18]. Our findings corroborate this bias and show induction of heat shock proteins and immediate-early response genes in warm-dissociated libraries when compared to cold-dissociated libraries. Cold-dissociated libraries can serve as a baseline in this case, since low temperature should minimise new transcription [18]. Our results further indicate that cell populations prone to these transcriptional changes are also depleted from the samples, with podocytes being the extreme example of a cell type practically lost in warm-dissociated libraries. Over-expression of stress response-related genes was also detected by bulk RNA-seq analysis of dissociated tissues, confirming that this artefact stems from the dissociation protocol rather than from microfluidic separation, single cell sequencing or data processing. These findings have important implications and suggest that data from samples digested at 37°C needs to be interpreted in light of this bias.

One possible drawback of cold tissue dissociation is lower efficiency of releasing hard-to-dissociate cell types. In our study, this may have affected cells of loop of Henle, which were detected at 2.52% in cold- and at 4.99% in warm-dissociated samples. However, both protocols dramatically under-estimated abundance of this second most populous kidney cell type. While one possible explanation could be incomplete tissue dissociation in both cases, deconvolution of bulk RNA-seq profiles of single-cell suspensions indicated that cells might be lost during cell encapsulation on the microfluidic device.

Two recent studies have shown that cryopreservation generated comparable data to that of fresh cells for cell lines and immune cells, and also for complex tissues cryopreserved prior to single-cell separation [22, 23]. Here, however, we report that cryopreservation of single-cell suspensions of dissociated mouse kidneys resulted in depletion of epithelial cell types. This artefact was reproducible across two mouse strains, both sexes, and two 10x chemistry versions. However, we observed a higher fraction of recovered PT cells (7.65% vs. 0.57%) in the repeated experiment, which might be explained by either sex or strain differences, or higher sensitivity of 10x v3 chemistry. Together with the depletion of PT cells, we observed reduced contamination of other cells with highly expressed PT transcripts, which indicates that PT cells might be lost in the thawing and resuspension. A possible explanation for the differences from previous reports is the proportion of serum used in the freezing media. 10x Genomics recommends 40% FBS (10x Genomics, CG00039, Rev D), whereas the other studies used either 90% FBS (peripheral blood, minced tissues, cell lines and immune cells) or 10% FBS (cell lines) [22, 23]. Notably, despite loading similar numbers of fresh, methanol-fixed and cryopreserved cells, the number of the usable cells observed in the cryopreserved samples was only ∼30% of the others. This raises the possibilities that the missing PT cells may be present but are failing to make it into the microfluidics, failing to lyse, or are so badly damaged that there is insufficient RNA remaining to generate a usable library. In contrast to cryopreservation, cell composition of methanol-fixed suspensions resembled that of freshly profiled libraries. Similarly to previous studies, we observed ambient RNA contamination with highly abundant transcripts suggesting cell damage by methanol fixation [22].

Studies comparing scRNA-seq and snRNA-seq reported that, although the two technologies profile different RNA fractions, both detect sufficient genes and allow adequate representation of cell populations [24–26]. In this work, one of the most notable differences between single-cell and single-nuclei experiments was the low detection rate of immune cells, in particular the failure to detect T, B, or NK cells in any of the snRNA-seq libraries. The depletion of lymphocytes was also observed in the Wu *et al.* [26] dataset (commented upon by O’Sullivan *et al.* [35]). Notably Slyper *et al.* [36] also observe much lower fractions of T, B, and NK cells in matched snRNA-seq and scRNA-seq datasets from adjacent pieces of a metastatic breast cancer and a neuroblastoma.

As Wu *et al.* have suggested, although these differences might indicate under-estimation of immune cells by snRNA-seq, another plausible explanation is that immune cell content is inflated in single-cell experiments as other cell types may be underrepresented due to incomplete dissociation. A major hurdle to determining which explanation is correct is the lack of a “ground truth” for cell composition of mouse kidneys. Clark *et al*. recently reported cell frequency estimates based on quantitative renal anatomy. However, these were restricted to renal epithelial cells [32]. Based on these estimates, some cell types, such as podocytes, appear to be better represented in scRNA-seq, whereas others, such as loop of Henle cells, were captured more effectively by snRNA-seq. We also attempted to use computational deconvolution of bulk RNA-seq of intact kidneys to infer its cell composition. However, the approach is sensitive to the input marker gene list used and may overlook rare and novel cell types. In addition, cell abundance estimates from bulk data would be influenced by both cell number and relative mRNA content of each cell. We will continue to search for approaches to better define the “ground truth” for cell composition.

## Conclusions

Our comparison of two tissue dissociation protocols revealed better performance of the cold dissociation protocol, while traditional digestion at 37°C introduced artifactual changes in sensitive cell populations affecting both representation of cell types and their transcriptional profiles. We also found that profiling of fresh single-cell suspensions is preferred; however, if immediate sample processing is challenging, methanol fixation gives satisfactory results introducing moderate cell damage. In contrast, cryopreservation of dissociated cells induces stress response and results in loss of the main epithelial cell type in kidney. Finally, we highlight differences in cell type composition between scRNA-seq and snRNA-seq libraries. Both approaches appear to have specific biases, thus when possible, studies would benefit from applying both for the same tissue.

## Methods

### Mice

Acknowledging the principles of 3Rs (Replacement, Reduction and Refinement) all kidneys used in this study were from mice that were euthanised by cervical dislocation as parts of other ongoing ethically approved experiments. In the first series of experiments, comparing cold and warm tissue dissociation and two preservation protocols, male AFAPIL.1DEL C57BL/6J mice from the same litter were used. These mice were 19 weeks old when euthanised and had no exposure to any experimental procedures. For the subsequent experiments, comparing cold-dissociated scRNA-seq to single-nuclei isolation protocols, we used untreated 18-week-old male Balb/c mice from the same litter or untreated 15-week-old female wild type Balb/c mice that were previously used as breeders, as specified in **Supplementary Table 17**.

### Kidney harvesting

Mice were euthanised and their kidneys were dissected and placed into a 1.5mL tube containing 1mL of ice-cold PBS. The capsules were then removed on ice and the samples processed as detailed below.

### Warm tissue dissociation

Kidneys were dissociated using the Multi-tissue dissociation kit 2 from Miltenyi Biotec [130-110-203] as per manufacturers’ instruction, with minor variations. Once the weight of the kidney was determined, the kidney was quartered and placed into a gentleMACS C-tube [Miltenyi Biotech; 130-096-334] containing the enzyme mix described in the kit’s protocol. The tube was centrifuged briefly, then placed onto the gentleMACS octo dissociator (Miltenyi Biotech), and the 37C_Multi_E program was run after attaching the heating elements. Following completion of the program, the tube was briefly centrifuged.

The homogenate was filtered through a 70µm cell strainer [Greiner; 54207] into a 50mL centrifuge tube [Greiner; 227270], the strainer was then rinsed with 15mL of PBS. The cell suspension was centrifuged at 400g for 10 minutes; once complete, the supernatant was removed and the pellet was resuspended in 5mL of PBS+0.04% BSA [Sigma; A7638]. The cell suspension was then filtered through a 40µm strainer [Greiner; 542040], which was subsequently rinsed with 2mL of PBS+0.04% BSA. The cells were again centrifuged at 400g for 10 minutes. The supernatant was then removed and the pellet was resuspended in 5mL of PBS+0.04% BSA. Cell count and viability was estimated using the Countess II FL (ThermoFisher) and the ReadyProbes Blue/Red kit [Invitrogen; R37610]. The cells were then diluted to 700 cells/µL and were immediately loaded onto a 10x chip A and processed on the 10x Chromium controller. The remaining cells were then either methanol fixed or cryopreserved.

### Cold tissue dissociation

Kidneys were dissociated using a modified version of the published protocol described in [18]. Based on the weight, in a pre-cooled Miltenyi C-tube, a protease solution (5mM CaCl2 [Invitrogen; AM9530G], 10mg/mL *B. Licheniformis* protease [Sigma; P5380], 125U/mL DNase I [Sigma; D5025], 1xDPBS) was prepared for each kidney.

The kidneys were then minced on ice into a smooth paste using a scalpel. The minced kidney was transferred into 4-6mL of the protease solution (dependent on weight) and triturated using a 1mL pipette for 15 seconds every two minutes for a total of eight minutes.

Following trituration, the C-tubes were placed onto a Miltenyi gentleMACS octo dissociator in a cool room (4°C), and the m_brain_03 program was run twice in succession. Once complete, the samples were triturated for 15 seconds every two minutes on ice for an additional 16 minutes using a 1mL pipette. 10µL of each sample was then loaded into a haemocytometer to assess whether tissue dissociation was complete. The dissociated cells were transferred to a 15mL centrifuge tube and 3mL of ice-cold PBS+10%FBS [Gibco; A3160401] was added.

The cell suspension was centrifuged at 1200g for five minutes at 4°C. The supernatant was removed and the pellet was resuspended in 2mL of PBS+10%FBS. The cells were then filtered through a 70µm cell strainer, which was subsequently rinsed with 2mL of PBS+0.01% BSA. The cells were then centrifuged again at 1200g for five minutes at 4°C followed by removal of the supernatant and resuspension of the pellet in 5mL of PBS+0.01%BSA. The cells were then filtered through a 40µm cell strainer, which was subsequently rinsed with 2mL of PBS+0.01% BSA. The cells were again centrifuged at 1200g for five minutes at 4°C followed by removal of the supernatant and resuspension of the cells in 5mL of PBS+0.04%BSA. The cells were counted and checked for viability using the ReadyProbes Blue/Red Kit on the Countess II FL. The cells were further diluted to a concentration of 700 cells/µL with PBS/0.04%BSA and loaded directly onto a 10x chip (A/B depending on experiment) and isolated using the 10x Chromium controller. The remaining cells were either methanol fixed or cryopreserved.

### Methanol fixation

#### Fixing

The methanol-fixation protocol was based on [37]. After tissue dissociation, the cells were concentrated to approximately 5x10^6^ cells/mL by centrifuging at 1000g for 10 minutes. 200µL of the cell suspensions were aliquoted into 2mL cryovials resting on ice. 800µL of 100% methanol [Sigma; 494437] (chilled at -20°C) was then added dropwise to each sample while gently stirring the cells to prevent clumping. The cryovials were stored at -20°C for 30 minutes, then directly transferred to -80°C (no gradient cooling).

#### Rehydrating

Cryovials of methanol fixed cells were removed from -80°C and placed on ice to equilibrate to 4°C (approximately 10 minutes). The cells were then transferred to a 1.5mL centrifuge tube and centrifuged at 1000g for five minutes at 4°C. The supernatant was discarded and the pellet was resuspended in a small volume of SSC cocktail (3xSSC [Sigma; S0902], 0.04% BSA, 40mM DTT [Sigma; 43816], 0.5U/mL RNasin plus [Promega; N2615]) to reach a concentration of approximately 2000 cells/µL. The cells were then filtered through a pre-wetted (with 1mL of nuclease-free water) 40µm pluristrainer mini filter [PluriSelect; 43-10040]. The cells were counted using the ReadyProbes Blue/Red Kit on the Countess II, then adjusted to 2000 cells/µL based on the count. The cells were loaded onto a 10x chip (A/B depending on version used) at a volume that dilutes the SSC to 0.125x to prevent reverse transcription inhibition.

### Cryopreservation

#### Freezing

After tissue dissociation, the cells were centrifuged at 400g for 10 minutes (1200g for five minutes at 4°C for the repeated experiment), then resuspended in freezing media (50% FBS, 40% RPMI-1640 [Gibco; 11875093], 10% DMSO [Sigma; D4540]) to achieve a concentration of 1x10^6^ cells/mL. 1mL of the cell suspension was aliquoted into 2mL cryovials, then placed into an isopropanol freezing container (Mr. Frosty) and stored at -80°C overnight. The following day, the cells were transferred to liquid nitrogen storage.

#### Thawing

The samples were removed from -80°C and immediately placed into a 37°C waterbath for 2-3 minutes to rapidly thaw. The cells were then mixed using a 1mL pipette with a wide-bore tip and the entire volume transferred to a 15mL centrifuge tube [Greiner; 188261]. The cryovial was then rinsed twice with RMPI+10%FBS (rinse media); each time the 1mL of media was added to the 15mL centrifuge in a dropwise manner while gently shaking the tube. 7mL of rinse media was added to the centrifuge tube using a serological pipette – the first 4mL was added dropwise while gently shaking the tube, and the following 3mL added down the side of the tube over two seconds. The tube was then inverted to mix.

The cells were centrifuged at 300g for five minutes. Once completed, the supernatant was removed (leaving 1mL), placed into another 15mL centrifuge tube and centrifuged at 400g for five minutes. The supernatant was discarded (leaving 1mL). The pellet from the supernatant was then resuspended, combined with the pellet in the initial centrifuge tube and mixed. 2mL of PBS+0.04% BSA was added to the centrifuge tube and shaken gently to mix. The cells were then centrifuged again at 400g for five minutes. The supernatant was discarded leaving 0.5mL behind. 0.5mL of PBS+0.04% BSA was added to the cells and gently pipette-mixed 10-15 times to fully resuspend. The cells were then filtered through a pre-wetted (with 1mL of PBS+0.04% BSA) 40µm pluristrainer mini filter. A 20µL aliquot of the cells was used to obtain an estimate of cell count and viability using the ReadyProbes Blue/Red Kit on the Countess II FL. Based on the count, the cells were diluted to a concentration of 700 cells/µL. The cells were then loaded onto a 10x chip (A/B depending on version) and immediately processed on the 10x Chromium controller.

For the repeated experiment, the above method was altered: Rather than a 300g spin followed by two 400g spins, two 1200g spins were performed, omitting the second centrifugation step.

### Flash freezing of whole kidney

Following the removal of the renal capsule, the kidney was placed into an isopentane [Sigma; 320404] bath resting on dry ice for five minutes. The temperature of the bath was maintained between -30°C and -40°C. Once frozen, the kidney was placed into a pre-cooled (on dry ice) cryovial and then buried in dry ice. The process was repeated for all designated kidneys. The flash-frozen kidneys were then transferred to a -80°C freezer for storage.

### Single nuclei isolation

#### SN_FANS_3x500g

This method is an adaptation of the Frankenstein protocol [38] and the 10x demonstrated protocol [39].

The kidneys were removed from -80°C and immediately placed on ice. Each kidney was then transferred to a 1.5mL tube containing 300µL of chilled lysis buffer (10mM Tris-HCl [Invitrogen; AM9856], 3mM MgCl2 [Invitrogen; AM9530G], 10mM NaCl [Sigma; 71386], 0.005% Nonidet P40 substitute [Roche; 11754599001], 0.2U/mL RNasin plus) and incubated on ice for two minutes. The tissue was then completely homogenised using a pellet pestle [Fisherbrand; FSB12-141-364] using up and down strokes without twisting. 1.2mL of chilled lysis buffer was added to the tube and pipette-mixed (wide-bore). The full volume was then transferred to a pre-cooled 2mL tube. The homogenate was incubated on ice for five minutes and mixed with a wide-bore tip every 1.5 minutes.

Following the incubation, 500µL of the lysis buffer was added to the homogenate, which was subsequently pipette-mixed and split equally into four 2mL tubes. 1mL of chilled lysis buffer was added to each tube and pipette-mixed using a wide-bore tip. The four tubes were incubated for a further five minutes on ice, mixing with a wide-bore tip every 1.5 minutes. The homogenate from the four tubes was then filtered through a 40µm strainer into a pre-cooled 50ml centrifuge tube. Following this, the sample was split again into four 2mL tubes resting on ice.

The samples were centrifuged at 500g for five minutes at 4°C. The supernatant was removed leaving 50µL in the tube. 1.5mL of lysis buffer was then added to two of the tubes and the pellet resuspended by mixing with a pipette. This resulted in two tubes containing 1.5mL resuspended nuclei in lysis buffer, and two tubes containing a nuclei pellet in 50µL of lysis buffer. The resuspended nuclei in one tube was then combined with the nuclei pellet of another, resulting in two tubes containing resuspended nuclei in lysis buffer.

The nuclei were centrifuged again at 500g for five minutes at 4°C. The supernatant was removed completely and discarded. 500µL of nuclei wash buffer (1xDPBS, 1% BSA, 0.2U/mL RNasin plus) was added to the tube containing the pellet and left to incubate without resuspending for five minutes. Following incubation, an additional 1mL of nuclei wash buffer was added, and the nuclei were resuspended by gently mixing with a pipette. The nuclei were again centrifuged at 500g for five minutes at 4°C, followed by discarding of the supernatant. The pellets were resuspended in 1.4mL of nuclei wash buffer, then transferred into a pre-cooled 1.5mL tube. Another 500g centrifugation step for five minutes at 4°C was performed. The supernatant was then discarded, and the nuclei pellet was resuspended in 1mL of nuclei wash buffer.

The nuclei were then filtered through a 40µm pluristrainer mini filter. 200µL of the filtered nuclei suspension was transferred into a 0.5mL tube and set aside to be used as the unstained control for sorting. To the remaining 800µL, 8µL of DAPI (10µg/mL) [ThermoScientific;. 62248] was added, and the nuclei were mixed with a pipette. A quality control step was performed by viewing the nuclei under a fluorescence microscope on a haemocytometer to check nuclei shape and count.

A BD Influx Cell Sorter was then used to sort 100,000 DAPI-positive events using a 70µm nozzle and a pressure of 22 psi (as per gating strategy, **Supplementary Fig. 10**). The post-sort nuclei concentration and quality were then checked using a fluorescence microscope and haemocytometer. Nuclei were then loaded onto a 10x chip (A/B depending on version used) and processed immediately on the 10x Chromium controller.

#### SN_FANS_1x2000g

The flash-frozen kidneys were removed from -80°C and transferred to a 1.5mL tube containing 500µL of pre-chilled lysis buffer same recipe as previous protocol) and allowed to rest on ice for two minutes. Each kidney was then homogenised with a pellet pestle with 40 up and down strokes without twisting the pellet. The resulting homogenate was mixed with a pipette and transferred to pre-cooled 15mL centrifuge tube containing 2mL of lysis buffer. The homogenate was incubated for 12 minutes on ice with mixing every two minutes using a glass fire-polished silanised Pasteur pipette [Kimble; 63A54]. Once incubation was complete, 2.5mL of nuclei wash buffer (same recipe as previous protocol) was added to the homogenate. The remaining tissue fragments were completely dissociated by repeated trituration of the homogenate using the glass Pasteur pipette.

The homogenate was then filtered through a 30µm MACS Smart Strainer [Miltenyi Biotech; 130-098-458] into a new 15mL centrifuge tube. The nuclei were centrifuged at 2000g for five minutes at 4°C. The supernatant was removed and the nuclei pellet was resuspended in 1mL of nuclei wash buffer. 200µL was aliquoted into a 0.5mL tube to be used as an unstained control for sorting. 8µL of DAPI (10µg/mL) was added to the remaining 800µL of nuclei. Quality and quantity of the nuclei was checked using a fluorescence microscope prior to sorting. Sorting and post sorting QC was performed in the same manner as for the SN_FANS_3x500g protocol. Nuclei were then loaded onto a 10x chip B and processed immediately on the 10x Chromium controller.

#### SN_sucrose

Kidneys were removed from -80°C and transferred to a 1.5mL tube containing 500µL of pre-chilled lysis buffer II (same recipe as previous protocols, with 125U/mL of DNase I added) and allowed to rest on ice for two minutes. Each kidney was then homogenised using a pellet pestle with 40 up and down strokes without twisting. The homogenate was transferred to a 15mL centrifuge tube containing 2mL of lysis buffer II and incubated for 12 minutes on ice with mixing every two minutes using a glass fire-polished silanised Pasteur pipette. Following the incubation, 2.5mL of nuclei wash buffer II (1xDPBS+2%BSA) was added to the homogenate. Remaining tissue clumps were dissociated by repeated trituration of the homogenate using the glass Pasteur pipette.

The homogenate was then filtered through a 30µm MACS Smart Strainer into a new 15mL centrifuge tube. Subsequently, the homogenate was centrifuged at 2000g for five minutes at 4°C. The supernatant was removed, and the pellet was resuspended in 510µL of nuclei wash buffer II. 10µL of the suspension was transferred to a 1.5mL tube and placed on ice for use in nuclei recovery calculations. 900µL of 1.8M sucrose solution [Sigma; NUC201] was added to the remaining 500µL of nuclei suspension and homogenised by mixing with a pipette. 3.6mL of 1.3M sucrose solution [Sigma; NUC201] was added to a 5mL tube. The nuclei/sucrose homogenate was then gently layered on top of the 1.3M sucrose solution.

The 5mL tube containing the sucrose solutions and nuclei was then centrifuged at 3000g for 10 minutes at 4°C. Once centrifugation was complete, the sucrose phase containing debris was soaked up using a Kimwipe wrapped around a pellet pestle. The remaining supernatant was removed and discarded using a pipette. The nuclei pellet was then resuspended in 5mL of wash buffer II, of which 10µL was transferred to a 1.5mL tube to assess nuclei recovery.

To the 10µL of nuclei suspension removed prior to the sucrose gradient, 980µL of wash buffer II and 10µL of DAPI (10µg/mL) was added. To the 10µL of nuclei suspension removed after the sucrose gradient, 89µL of wash buffer II and 1µL of DAPI (10µg/mL) was added. The yield from the pre- and post-sucrose aliquots were compared to assess nuclei recovery after filtration through the gradient. The post-sucrose count was used to dilute the nuclei to a concentration of 700 nuclei/µL, which was immediately loaded onto a 10x Chip B and processed with the 10x Chromium controller.

### Single cell RNA-seq library preparation

All single cell libraries were constructed in biological triplicate using the 10x Chromium 3’ workflow as per manufacturers’ directions. In the first series of experiments, comparing cold and warm tissue dissociation and two preservation protocols, version two chemistry was used. For single-cell versus single-nuclei comparisons, versions two and three were used as indicated in **Fig. 4A**. All experiments and conditions aimed for a capture of approximately 9000 cells, except for methanol fixed samples. Due to the reverse transcription inhibition of 3x SSC, the sample had to be loaded at a concentration of 0.125x SSC, resulting in an approximate cell capture of 4000-5000 cells.

### Bulk RNA-seq library preparation

For the undissociated samples, total RNA was extracted from flash-frozen kidneys using the Nucleospin RNA Midi kit [Macherey Nagel; 740962.20] as per manufacturers’ directions. For the dissociated samples, total RNA was extracted from the remaining cells from each of the tissue dissociation protocols. RNA was assessed for quantity and quality using the TapeStation 4200 RNA ScreenTape kit [Agilent; 5067-5576], which showed all RNA used had a RIN of>8. Bulk RNA-seq was performed using the NEBNext Ultra II RNA Library Kit for Illumina [NEB; E7760] and either NEBNext rRNA Depletion Kit (Human/Mouse/Rat) [NEB; E6310] or NEBNext Poly(A) mRNA isolation module [NEB; E7490] as described in the manufacturers’ protocol, with 100ng of total RNA as input.

### Sequencing

All libraries were quantified with qPCR using the NEBnext Library Quant Kit for Illumina and checked for fragment size using the TapeStation D1000 kit (Agilent). The libraries were pooled in equimolar concentration for a total pooled concentration of 2nM. 10x single cell libraries were sequenced using the Illumina NovaSeq 6000 and S2 flow cells (100 cycle kit) with a read one length of 26 cycles, and a read two length of 92 or 98 cycles for version two chemistry. Version three chemistry had a read one length of 28 cycles, and a read two length of 94 cycles. Bulk libraries were sequenced on the Illumina NovaSeq 6000 using SP flow cells (100 cycle kit) with read length of 150 for dissociated bulk in C57BL/6J mice, 51 for undissociated bulk in Balb/c male mice, and 60 for Balb/c female mice (**Supplementary Table 17**).

### Bulk RNA-seq data processing

BCL files were demultiplexed and converted into FASTQ using bcl2fastq utility of Illumina BaseSpace Sequence Hub. FastQC was used for read quality control (https://www.bioinformatics.babraham.ac.uk/projects/fastqc/). Adapters and low-quality bases were trimmed using Trim Galore with parameters *--paired --quality 5 --stringency 5 --length 20 --max_n 10* (https://www.bioinformatics.babraham.ac.uk/projects/trim_galore/). Reads matching to ribosomal DNA repeat sequence BK000964 (https://www.ncbi.nlm.nih.gov/nuccore/BK000964) and low complexity reads were removed with TagDust2 [40]. The remaining reads were mapped to GRCm38.84 version of mouse genome using STAR version 2.6.1a with default settings [41]. Picard MarkDuplicates tool was employed to identify duplicates (https://broadinstitute.github.io/picard/command-line-overview.html#MarkDuplicates). FeatureCounts was then used to derive gene count matrix [42]. Counts were normalised to gene length and then to library sizes using weighted trimmed mean of M-values (TMM) method in edgeR [27], to derive gene length corrected trimmed mean of M-values (GeTMM) as described in [43].

### scRNA-seq and snRNA-seq data processing

BCL files were demultiplexed and converted into FASTQ using bcl2fastq utility of Illumina BaseSpace Sequence Hub. scRNA-seq and snRNA-seq libraries were processed using Cell Ranger 2.1.1 with mm10-2.1.0 reference. Reads mapped to exons were used for scRNA-seq samples, whereas both intronic and exonic reads were counted for snRNA-seq. Custom pre-mRNA reference for snRNA-seq was built as described in https://support.10xgenomics.com/single-cell-gene-expression/software/pipelines/latest/advanced/references#premrna. Raw gene-barcode matrices from Cell Ranger output were used for downstream processing. Cells were distinguished from background noise using EmptyDrops [44]. Only genes detected in a minimum of 10 cells were retained; cells with 200-3000 genes and under 50% of mitochondrial reads were retained, as per Park et al. study [30]. Nuclei were additionally filtered to have at least 450 UMIs for v2 chemistry and 900 UMIs for v3 chemistry, and mitochondrial genes were removed. Outlier cells with high ratio of number of detected UMI to genes (>3 median absolute deviations from median) were removed using Scater [45]. Seurat v2 was used for sample integration (canonical correlation analysis), normalisation (dividing by the total counts, multiplying by 10,000 and natural-log transforming), scaling, clustering and differential expression analysis (Wilcoxon test) [34].

### Inferring cell identity

To infer cell identity for freshly profiled samples in the first series of experiments, we performed a reference-based annotation using scMatch [29] and refined cell labels based on marker gene expression in a two-step procedure described below (**Supplementary Fig. 1**).

#### Reference dataset

To construct the reference dataset for scMatch [29], we obtained gene counts and cell types reported in three single-cell (or single-nuclei) adult mouse kidney studies [26, 30, 31]. Counts were normalised to cell library size and averaged within each cell type to derive reference vectors (**Supplementary Fig. 1, Step 1**). The reference vectors were clustered using Spearman correlation coefficient and five vectors were removed as outliers. The remaining 66 vectors composed a reference dataset, available as **Supplementary Table 11**. With this reference dataset, we ran scMatch [29] (**Supplementary Fig. 1, Step 2**) using options *--testMethod s --keepZeros y* to label each individual cell with the closest cell type identity from the reference dataset.

#### Refining cell identities

We next refined scMatch-derived cell types based on gene expression. First, for each cell type, we calculated gene signatures as genes over-expressed in the given cell type when compared to all other cells (*FindMarkers* function of Seurat [34], minimum detection rate of 0.5, logFC threshold of 1 and FDR < 0.05 were used as thresholds; only cell types with at least 10 cells were considered; **Supplementary Fig. 1, Step 3**). Second, cell type gene signature scores were calculated for each cell and for each gene signature (*AddModuleScore* function of Seurat [34], genes attributed to signatures in more than two cell types were excluded; **Supplementary Fig. 1, Step 4**). Third, we used these scores to assign cell types to cells (**Supplementary Fig. 1, Step 5**). A cell type was assigned to a cell if the score for that cell type was the highest among all cell types, positive and significant with FDR < 0.05. Significance was determined in a Monte-Carlo procedure with 1000 randomly selected gene sets of the same size [46], correction for multiple hypothesis testing was performed using Benjamini-Hochberg procedure [47]. Cells without cell type annotation were manually explored to identify whether the corresponding cell type might be a novel one, absent from the reference.

#### Second iteration

Cell types inferred in our dataset were added to the reference dataset (**Supplementary Fig. 1, Step 6**), and annotation with scMatch and gene set signature scoring was repeated. Cells left unannotated at this stage were labelled as “unknown”. Cell type gene signatures are available in **Supplementary Table 12**.

This approach failed to identify cells of connecting tubule (CNT) and, instead, matched them to other similar cell types. To resolve this, annotation for cell types labelled as DCT, aLOH, CD_IC, CD_PC, CD_Trans was additionally refined as follows. These cells were extracted from the dataset and clustered separately. Candidate CNT cells were identified as a cluster over-expressing *Calb1* and *Klk1* genes [32, 48]. The cell type signature score procedure was then applied for this subset as described above.

Cell type labels assigned to each cell are available in **Supplementary Table 2**.

#### Preserved cells

Cells of preserved single-cell suspensions from the first series of experiments were annotated using the cell type gene signatures derived from the corresponding freshly profiled samples (**Supplementary Table 12**) and the gene set signature scoring procedure described above. Cell type labels assigned to each cell are available in **Supplementary Table 2**.

#### Subsequent experiments

In subsequent experiments, we used a combined reference dataset, which included the public data as well as data from freshly profiled cells generated in the first series of experiments (**Supplementary Table 13**, note that two cell types were excluded from the reference as outliers). Single-cell datasets were annotated using a single iteration of scMatch. For single-nucleus datasets we repeated the two-step annotation procedure described above. Cell type labels assigned to each cell or nucleus are available in **Supplementary Table 2**.

### Stress response score

Stress response score was calculated for 17 genes (*Fosb, Fos, Jun, Junb, Jund, Atf3, Egr1, Hspa1a, Hspa1b, Hsp90ab1, Hspa8, Hspb1, Ier3, Ier2, Btg1, Btg2, Dusp1*) for each cell using *AddModuleScore* function of Seurat version 2 [34]. The score represents an average expression levels of these genes on single-cell level, subtracted by the aggregated expression of control gene sets. All analysed genes were binned based on averaged expression, and the control genes were randomly selected from each bin. Significance was determined in a Monte-Carlo procedure with 1000 randomly selected sets of 17 genes [46], correction for multiple hypothesis testing was performed using Benjamini-Hochberg procedure [47].

### Cell cycle phase prediction

Cell cycle phases were inferred using *CellCycleScoring* function of Seurat version 2 [34] with the following genes: S-genes: *Atad2, Blm, Brip1, Casp8ap2, Ccne2, Cdc45, Cdc6, Cdca7, Chaf1b, Clspn, Dscc1, Dtl, E2f8, Exo1, Fen1, Gins2, Gmnn, Hells, Mcm2, Mcm4, Mcm5, Mcm6, Msh2, Nasp, Pcna, Pcna-ps2, Pola1, Pold3, Prim1, Rad51ap1, Rfc2, Rpa2, Rrm1, Rrm2, Slbp, Tipin, Tyms, Ubr7, Uhrf1, Ung, Usp1, Wdr76*; G2M-genes: *Anln, Anp32e, Aurka, Aurkb, Birc5, Bub1, Cbx5, Ccnb2, Cdc20, Cdc25c, Cdca2, Cdca3, Cdca8, Cdk1, Cenpa, Cenpe, Cenpf, Ckap2, Ckap2l, Ckap5, Cks1brt, Cks2, Ctcf, Dlgap5, Ect2, G2e3, Gas2l3, Gtse1, Hjurp, Hmgb2, Hmmr, Kif11, Kif20b, Kif23, Kif2c, Lbr, Mki67, Ncapd2, Ndc80, Nek2, Nuf2, Nusap1, Psrc1, Rangap1, Smc4, Tacc3, Tmpo, Top2a, Tpx2, Ttk, Tubb4b, Ube2c*. Note cells not annotated as S or G2M phase are by default labelled as G1 phase.

### Bulk RNA-seq deconvolution

BSEQ-sc was used for bulk expression deconvolution [33]. In the first series of experiments, marker genes for the deconvolution were calculated from scRNA-seq data, using only cold-dissociated samples to avoid the influence of the identified warm dissociation-related biases. We also excluded cells labelled as “Unknown” and “CD_Trans” from the calculation. For each of the remaining cell types, marker genes were calculated using Seurat function FindMarkers with the following thresholds: *logfc.threshold=1.5, min.pct = 0.5, only.pos = T*. Genes identified in more than one cell type were removed and the remaining genes were used for the deconvolution. The same set of genes was used to deconvolve all bulk RNA-seq libraries.

## Author Contributions

Conception and design: AF. Analysis and interpretation of data: ED with help from RH and TL. Writing, review and revision of the manuscript: ED and AF with input from all authors. DP and BS developed the SN_FANS_1x2000g protocol, OC developed the SN_sucrose protocol. BS and DP performed FANS. BG adapted the dissociation and SN_FANS_3x500g protocols. MJ, BG and LdK generated the single cell/nucleus and bulk libraries. MJ and LdK adapted the SN_sucrose protocol. MJ performed the sequencing for the libraries. Study supervision: AF.

## Data Availability Statement

Data that support the findings of this study have been deposited in GEO with the primary accession code GSE141115.

## Supplementary Tables

**Supplementary Table 1.** Genes differentially expressed between bulk RNA-seq profiles of cold- and warm-dissociated kidney single-cell suspensions (FDR < 0.05 and logFC > 2, edgeR exact test [27]); includes results of functional analysis with ToppGene [28] and Gene Ontology Biological Process for differentially expressed genes with higher expression in warm-dissociated kidneys.

**Supplementary Table 2.** Cell type labels assigned to cells and nuclei in this study. aLOH: ascending loop of Henle; CD_IC: intercalated cells of collecting duct; CD_IC_A: type A intercalated cells of collecting duct; CD_IC_B: type B intercalated cells of collecting duct; CD_PC: principal cells of collecting duct; CD_Trans: transitional cells; CNT: connecting tubule; DCT: distal convoluted tubule; dLOH: descending loop of Henle; MC: mesangial cells; MPH: macrophages; PT: proximal tubule.

**Supplementary Table 3.** Differentially expressed genes with higher expression in cell populations of warm-dissociated kidneys (Seurat Wilcoxon test [34] with thresholds of logFC = 0.5, minimum detection rate 0.5, FDR < 0.05); includes an incidence table indicating in how many cell types each gene was identified as differentially expressed and results of functional analysis with ToppGene [28] and Gene Ontology Biological Process for differentially expressed genes identified in at least one cell type.

**Supplementary Table 4.** Differentially expressed genes with higher expression in cell populations of cold-dissociated kidneys (Seurat Wilcoxon test [34] with thresholds of logFC = 0.5, minimum detection rate 0.5, FDR < 0.05); includes an incidence table indicating in how many cell types each gene was identified as differentially expressed.

**Supplementary Table 5.** Genes differentially expressed between cryopreserved and freshly profiled cold-dissociated kidney single-cell suspensions (Seurat Wilcoxon test [34] with thresholds of logFC = 1, minimum detection rate 0.5, FDR < 0.05), positive logFC indicates higher expression in cryopreserved samples; includes incidence tables indicating in how many cell types each gene was identified as differentially expressed.

**Supplementary Table 6.** Genes differentially expressed between cryopreserved and freshly profiled warm-dissociated kidney single-cell suspensions (Seurat Wilcoxon test [34] with thresholds of logFC = 1, minimum detection rate 0.5, FDR < 0.05), positive logFC indicates higher expression in cryopreserved samples; includes incidence tables indicating in how many cell types each gene was identified as differentially expressed.

**Supplementary Table 7.** Genes differentially expressed between methanol-fixed and freshly profiled cold-dissociated kidney single-cell suspensions (Seurat Wilcoxon test [34] with thresholds of logFC = 1, minimum detection rate 0.5, FDR < 0.05), positive logFC indicates higher expression in methanol-fixed samples; includes incidence tables indicating in how many cell types each gene was identified as differentially expressed.

**Supplementary Table 8.** Genes differentially expressed between methanol-fixed and freshly profiled warm-dissociated kidney single-cell suspensions (Seurat Wilcoxon test [34] with thresholds of logFC = 1, minimum detection rate 0.5, FDR < 0.05), positive logFC indicates higher expression in methanol-fixed samples; includes incidence tables indicating in how many cell types each gene was identified as differentially expressed.

**Supplementary Table 9.** Number of cells in each cell population across single-cell and single-nuclei experiments and BSEQ-sc estimates for bulk RNA-seq libraries.

**Supplementary Table 10.** Genes differentially expressed between single-cell and single-nuclei libraries in Balb/c male mice profiled with v2 10x chemistry (Seurat Wilcoxon test [34] with thresholds of logFC = 0.5, minimum detection rate 0.5, FDR < 0.05), positive logFC indicates higher expression in single-cell libraries; includes incidence tables indicating in how many cell types each gene was identified as differentially expressed.

**Supplementary Table 11.** Public reference dataset used for the first scMatch run.

**Supplementary Table 12.** Cell type gene signatures for refined annotation.

**Supplementary Table 13.** Extended reference dataset used for scMatch annotation.

**Supplementary Table 14.** Genes differentially expressed in each cell type between SN_FANS_1x2000g and SN_FANS_3x500g protocols (Seurat Wilcoxon test [34] with thresholds of logFC = 0.5, minimum detection rate 0.5, FDR < 0.05), positive logFC indicates higher expression in SN_FANS_1x2000g libraries; includes incidence tables indicating in how many cell types each gene was identified as differentially expressed.

**Supplementary Table 15.** Genes differentially expressed in each cell type between SN_FANS_1x2000g and SN_sucrose protocols (Seurat Wilcoxon test [34] with thresholds of logFC = 0.5, minimum detection rate 0.5, FDR < 0.05), positive logFC indicates higher expression in SN_FANS_1x2000g libraries; includes incidence tables indicating in how many cell types each gene was identified as differentially expressed.

**Supplementary Table 16.** Genes differentially expressed in each cell type between SN_sucrose and SN_FANS_3x500g protocols (Seurat Wilcoxon test [34] with thresholds of logFC = 0.5, minimum detection rate 0.5, FDR < 0.05), positive logFC indicates higher expression in SN_sucrose libraries; includes incidence tables indicating in how many cell types each gene was identified as differentially expressed.

**Supplementary Table 17**. Mice used and workflows tested. The experiments were performed in three batches separated in time. For each batch, we used kidneys from mice available at that time and the workflows were used with modifications as indicated in this table.

**Supplementary Table 18.** Genes differentially expressed between bulk RNA-seq profiles of intact kidneys and cold-dissociated single-cell suspensions; includes results of functional analysis with ToppGene [28] for genes with higher expression in intact kidneys.

## Supplementary Figures

**Supplementary Fig. 1.**
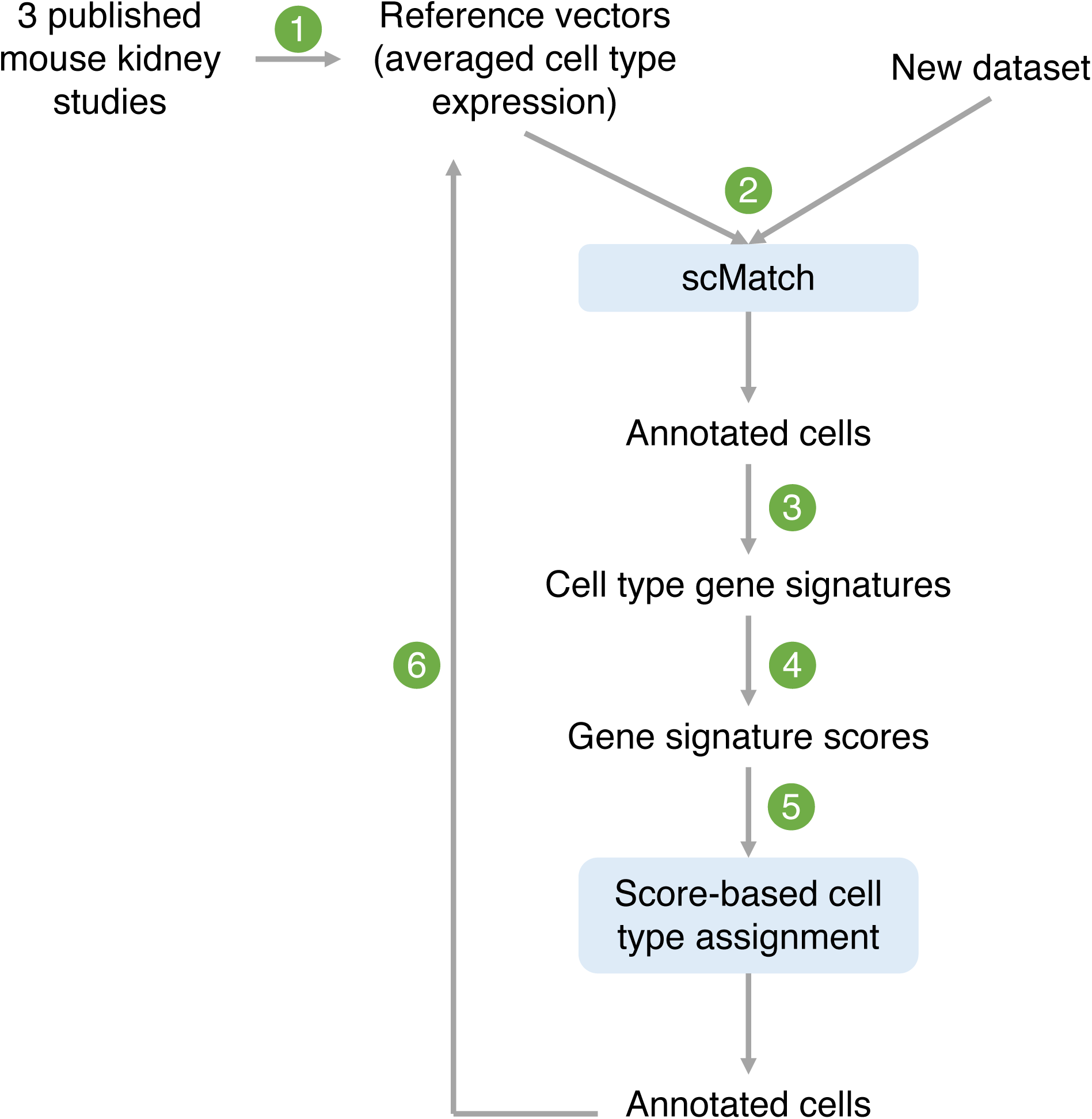
Cell annotation procedure. See **Methods** for more details.

**Supplementary Fig. 2.**
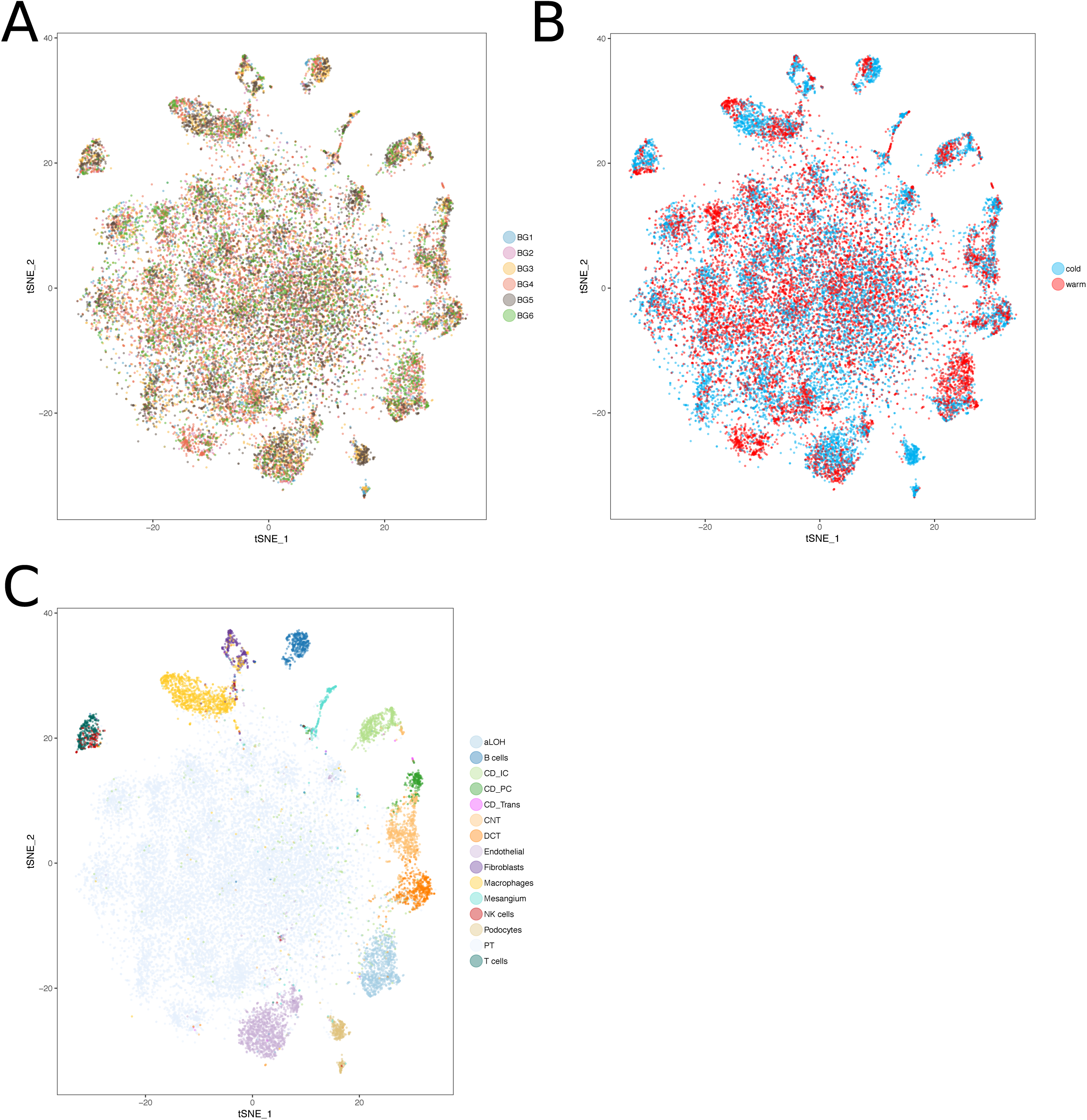
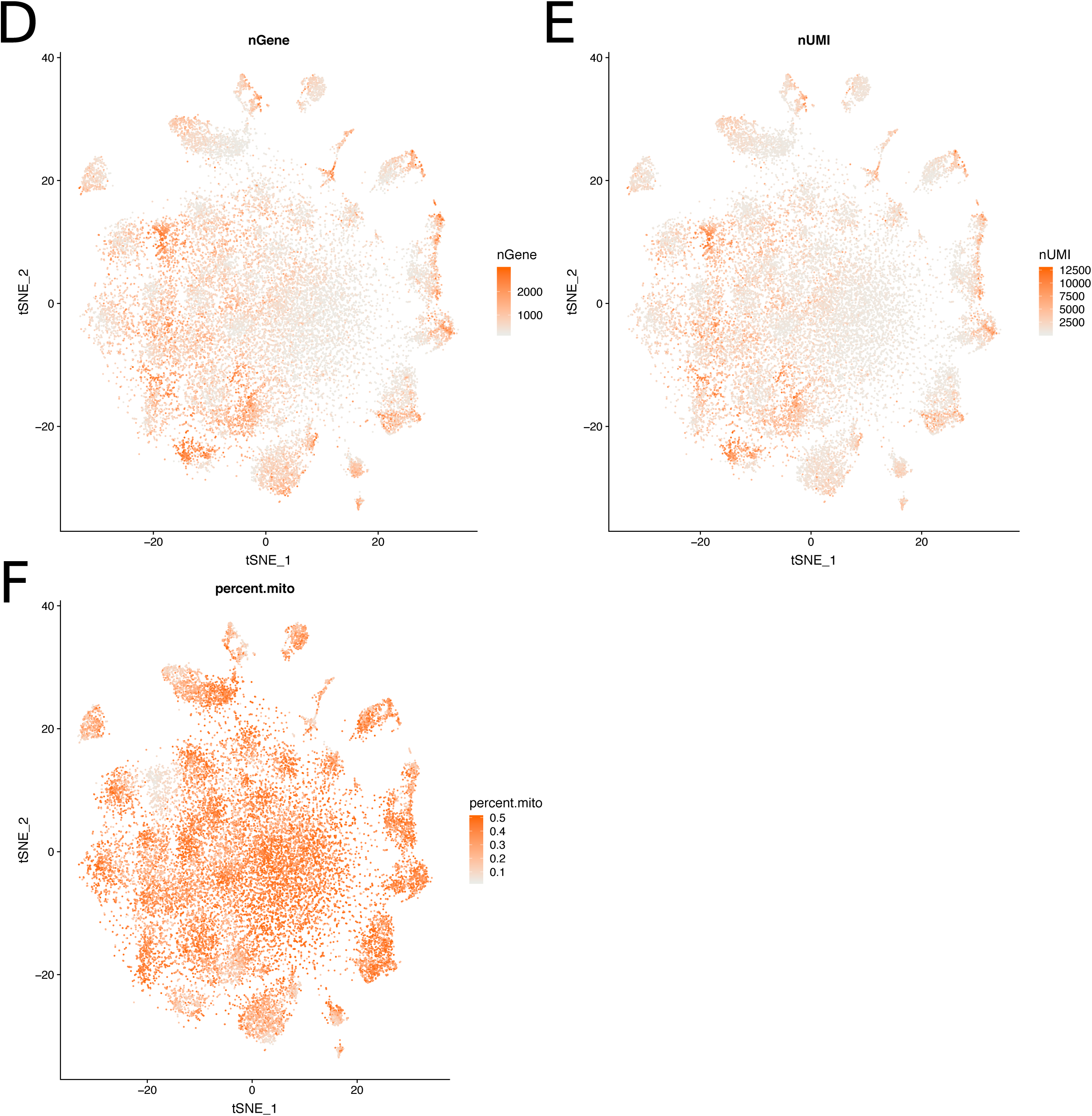
TSNE plots showing freshly profiled cells from cold- and warm-dissociated kidneys coloured by **a**) library, **b**) dissociation protocol, **c**) inferred cell type, **d**) number of detected genes, **e**) number of UMIs, **f**) fraction of reads mapped to mitochondrial genes. aLOH: ascending loop of Henle; CD_IC: intercalated cells of collecting duct; CD_PC: principal cells of collecting duct; CD_Trans: transitional cells of collecting duct; CNT: connecting tubule; DCT: distal convoluted tubule; PT: proximal tubule.

**Supplementary Fig. 3.**
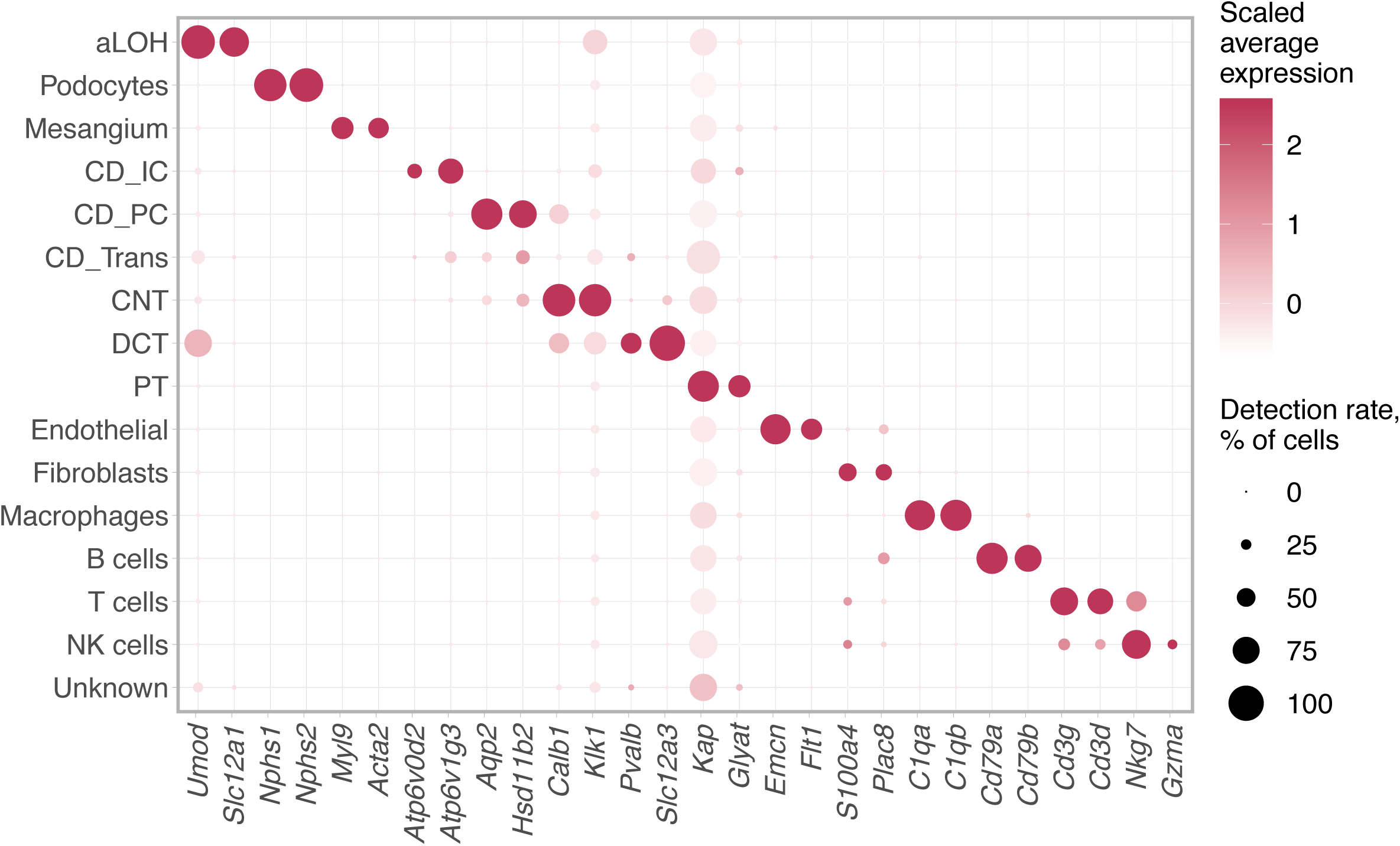
Expression and detection levels of selected marker genes. aLOH: ascending loop of Henle; CD_IC: intercalated cells of collecting duct; CD_PC: principal cells of collecting duct; CD_Trans: transitional cells of collecting duct; CNT: connecting tubule; DCT: distal convoluted tubule; PT: proximal tubule.

**Supplementary Fig. 4.**
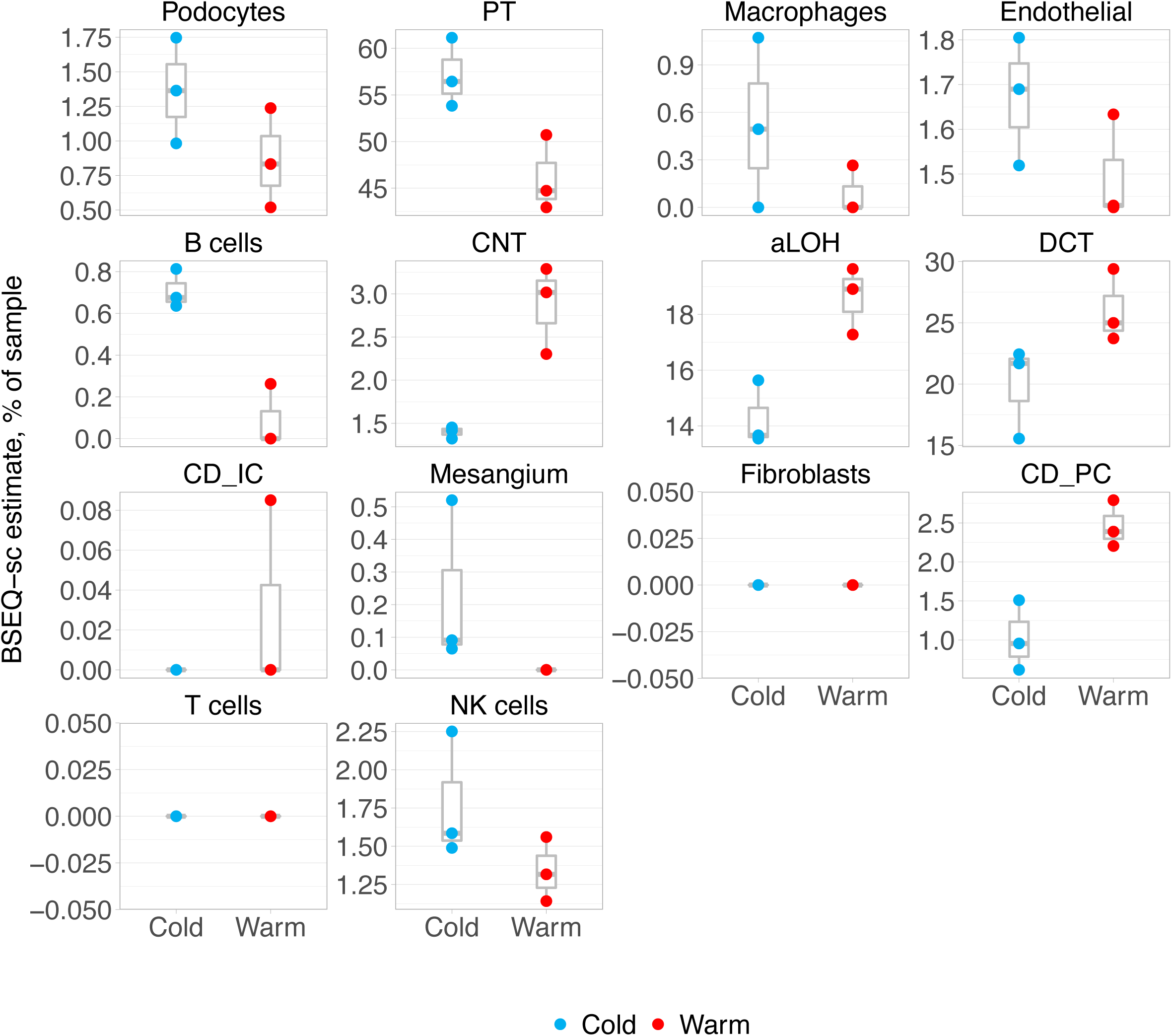
BSEQ-sc deconvolution of bulk RNA-seq profiles of cold- and warm-dissociated kidney single-cell suspensions. Three biological replicates are shown per condition. aLOH: ascending loop of Henle; CD_IC: intercalated cells of collecting duct; CD_PC: principal cells of collecting duct; CNT: connecting tubule; DCT: distal convoluted tubule; PT: proximal tubule.

**Supplementary Fig. 5.**
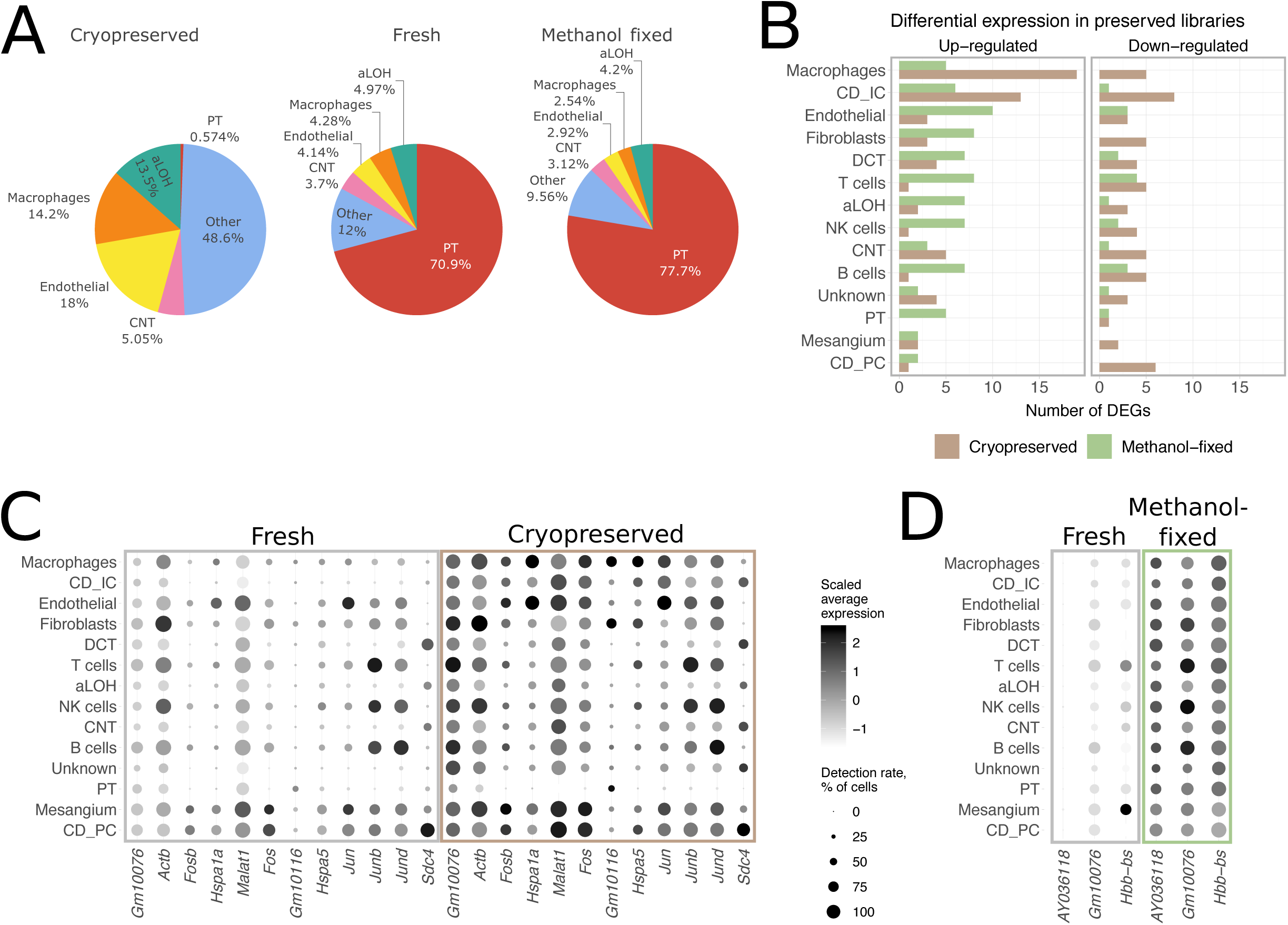
Cell preservation protocol performance in warm-dissociated samples. **a**) Cell type composition of fresh and preserved warm-dissociated samples. **b**) Number of DEGs detected between preserved and freshly profiled aliquots. Seurat Wilcoxon test [34] with logFC = 1, min detection rate 0.5, FDR < 0.05 as thresholds. **c**) Expression and detection rates of DEGs with higher expression in cryopreserved samples in at least two cell types. **d**) Expression and detection rates of DEGs with higher expression in methanol-fixed samples in at least nine cell types. aLOH: ascending loop of Henle; CD_IC: intercalated cells of collecting duct; CD_PC: principal cells of collecting duct; CD_Trans: transitional cells of collecting duct; CNT: connecting tubule; DCT: distal convoluted tubule; PT: proximal tubule.

**Supplementary Fig. 6.**
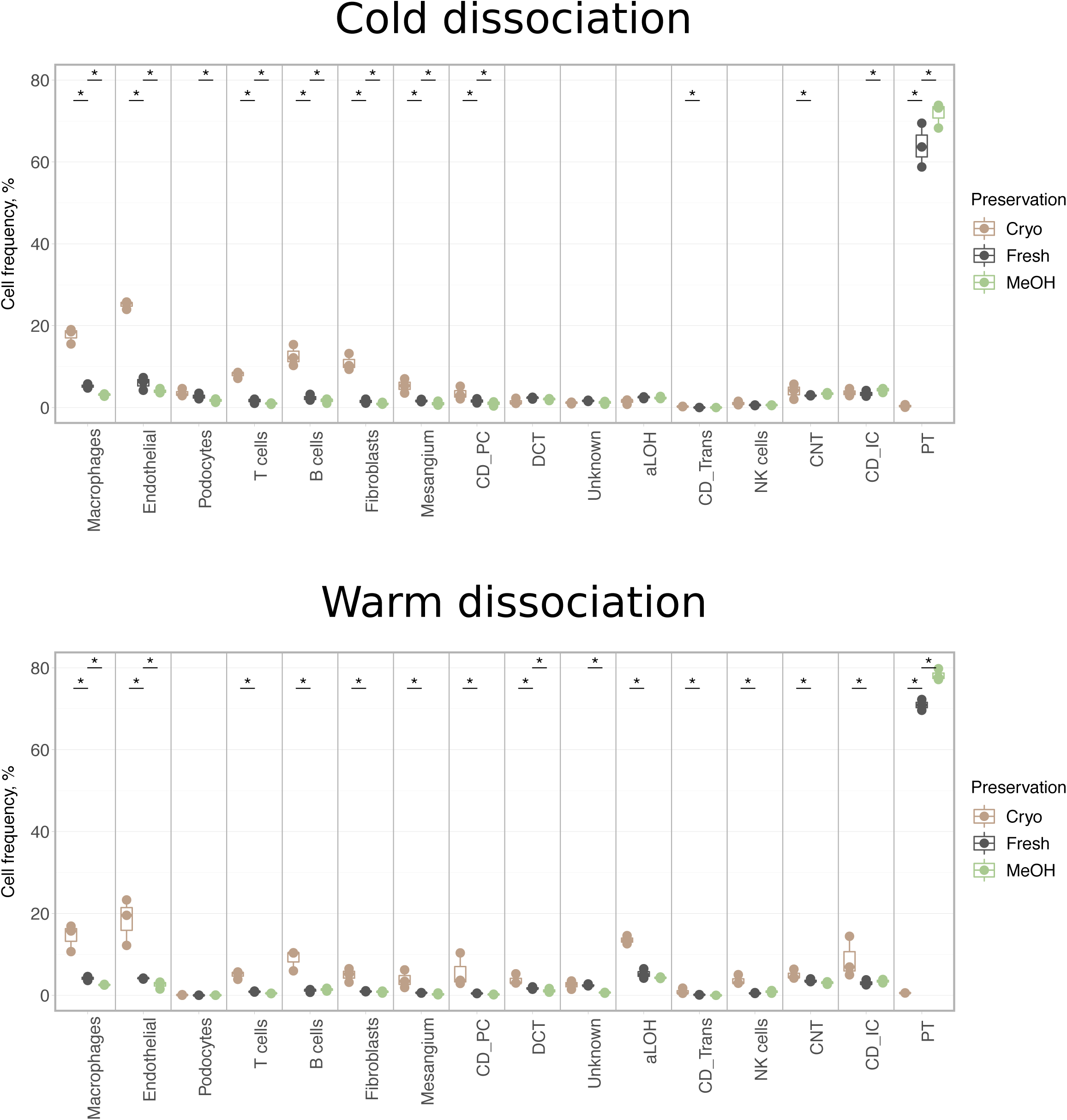
Cell type composition of fresh and preserved kidneys. Three biological replicates are shown per condition. Asterisks denote two-sided chi-square test p-value < 0.001. aLOH: ascending loop of Henle; CD_IC: intercalated cells of collecting duct; CD_PC: principal cells of collecting duct; CD_Trans: transitional cells of collecting duct; CNT: connecting tubule; DCT: distal convoluted tubule; PT: proximal tubule.

**Supplementary Fig. 7.**
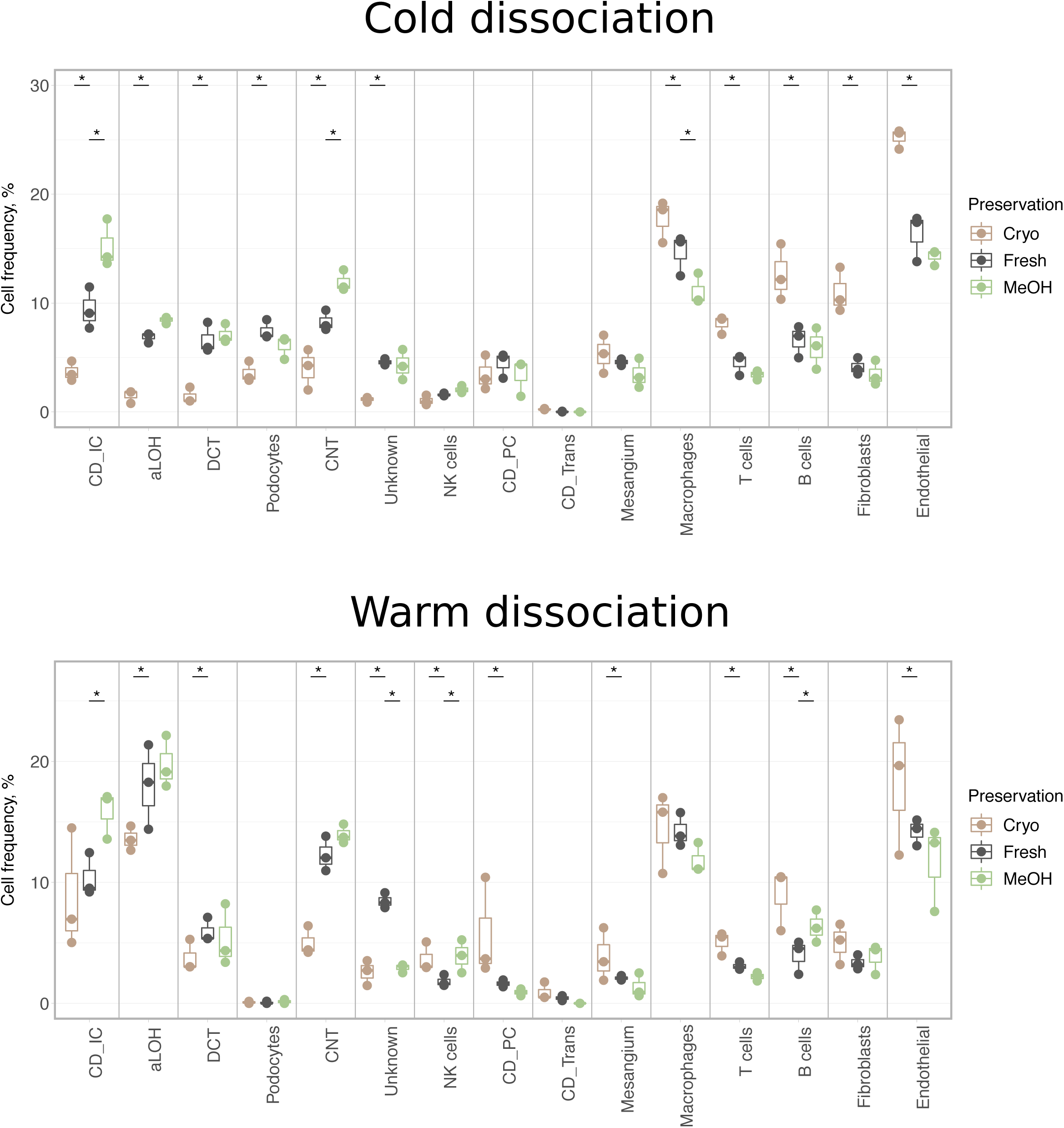
Cell type composition of non-proximal tubule cells in fresh and preserved kidneys. Three biological replicates are shown per condition. Asterisks denote two-sided chi-square test p-value < 0.001. aLOH: ascending loop of Henle; CD_IC: intercalated cells of collecting duct; CD_PC: principal cells of collecting duct; CD_Trans: transitional cells of collecting duct; CNT: connecting tubule; DCT: distal convoluted tubule.

**Supplementary Fig. 8.**
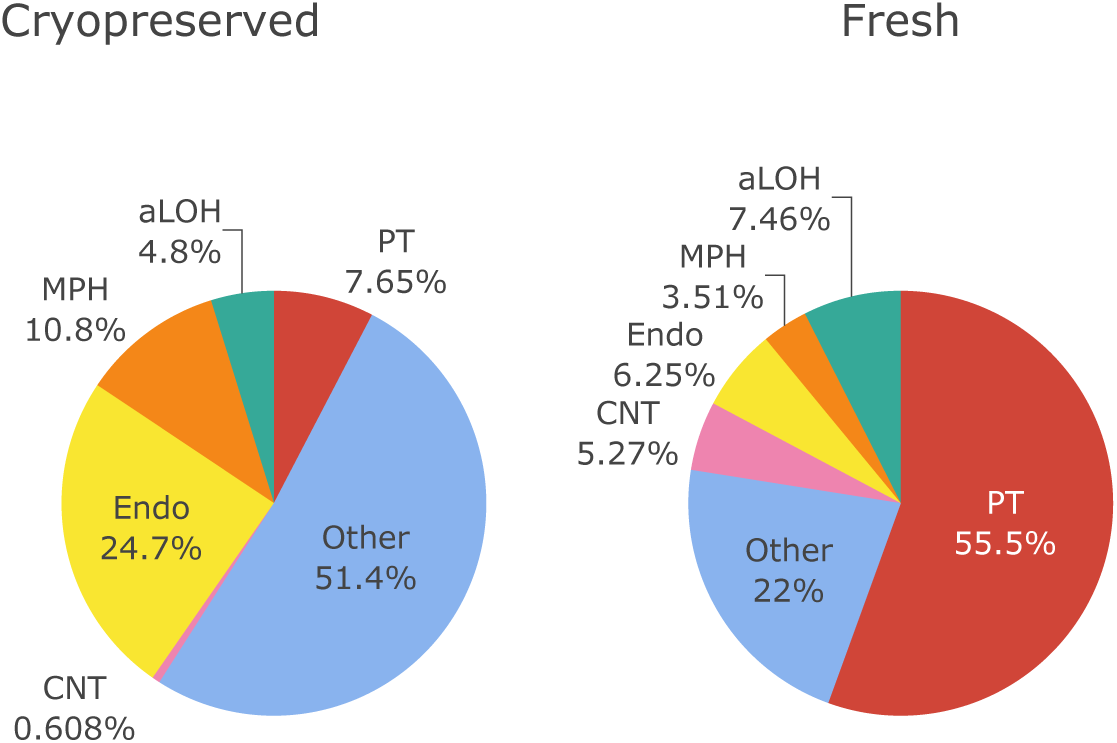
Cell type composition of fresh and cryopreserved cold-dissociated samples in the repeated experiment using Balb/c female mice, 10x v3 chemistry, 2 weeks storage, and 1200g spin. aLOH: ascending loop of Henle; MPH: macrophages; CNT: connecting tubule; PT: proximal tubule.

**Supplementary Fig. 9.**
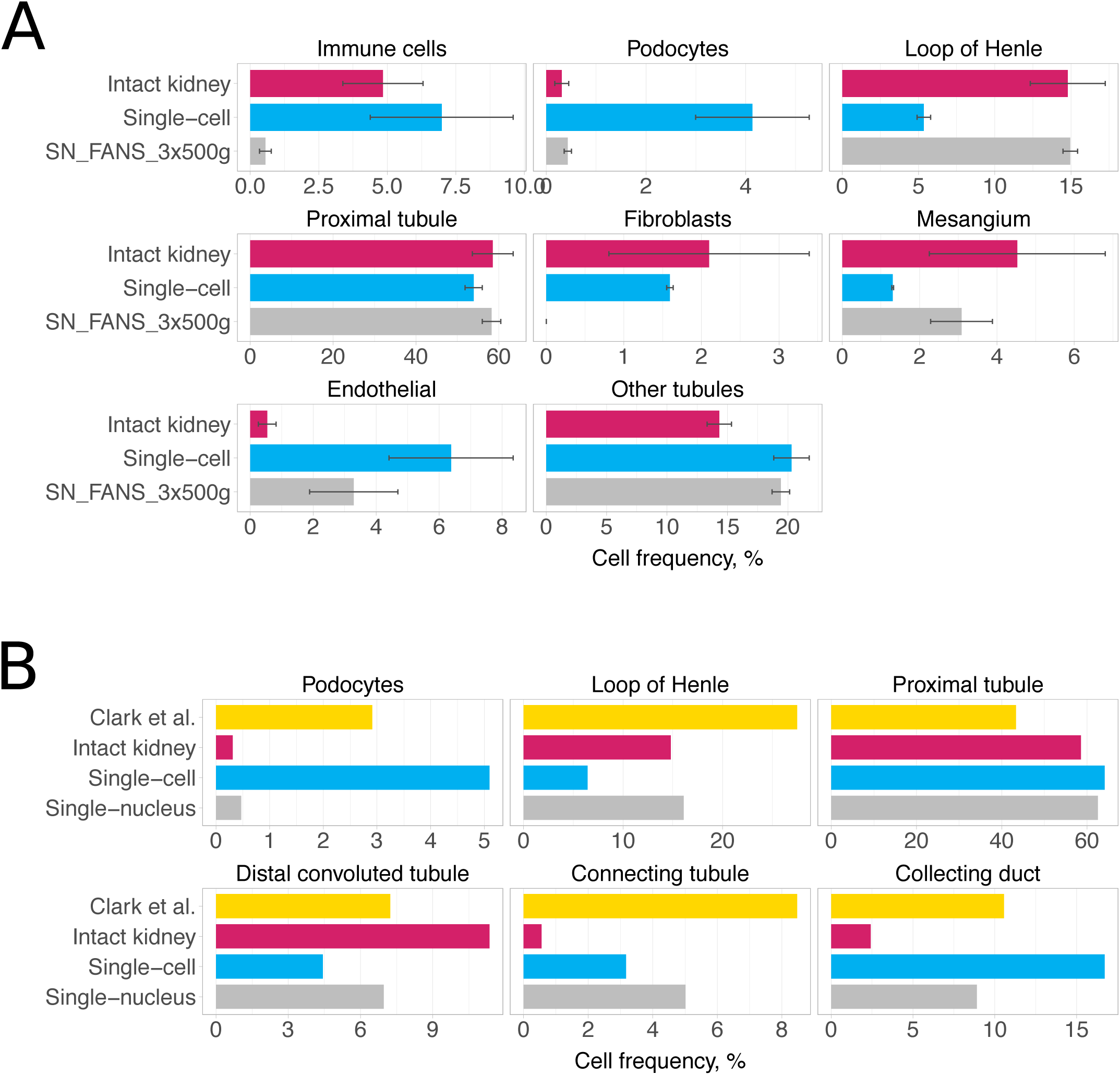
Comparison of single-cell and single-nucleus libraries in Balb/c male mice. **a**) Cell type composition for kidneys from Balb/c male mice. Average percentages for scRNA-seq libraries are shown in blue and for snRNA-seq libraries in grey. BSEQ-sc estimates are shown for bulk RNA-seq of intact kidneys. Error bars are standard error of mean. **b**) Abundance of renal epithelial cell types in Clark *et al*. study [32] in comparison to our data from Balb/c male mice.

**Supplementary Fig. 10.**
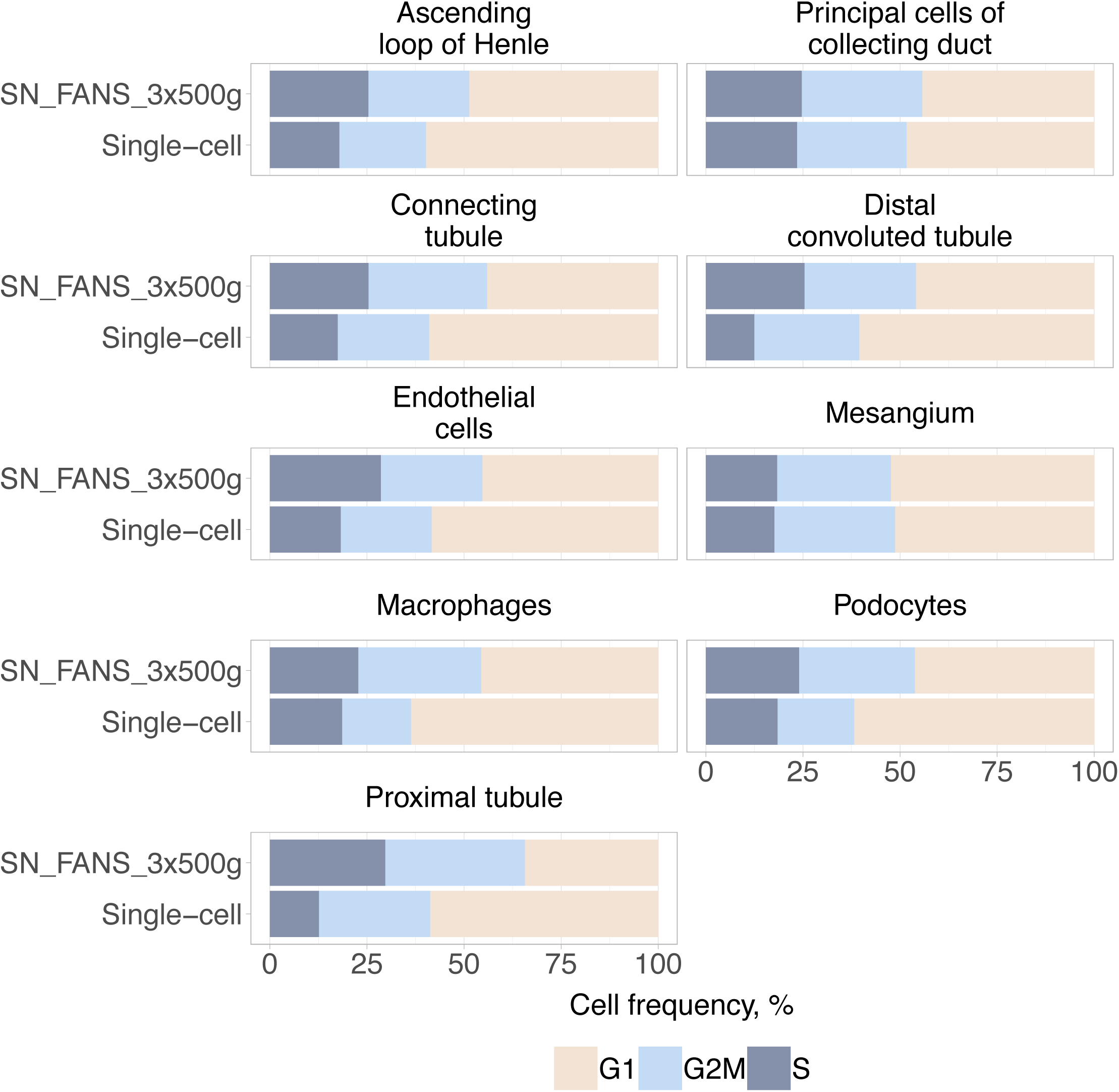
Cell cycle phases inferred in scRNA-seq and snRNA-seq libraries from Balb/c male mice.

**Supplementary Fig. 11.**
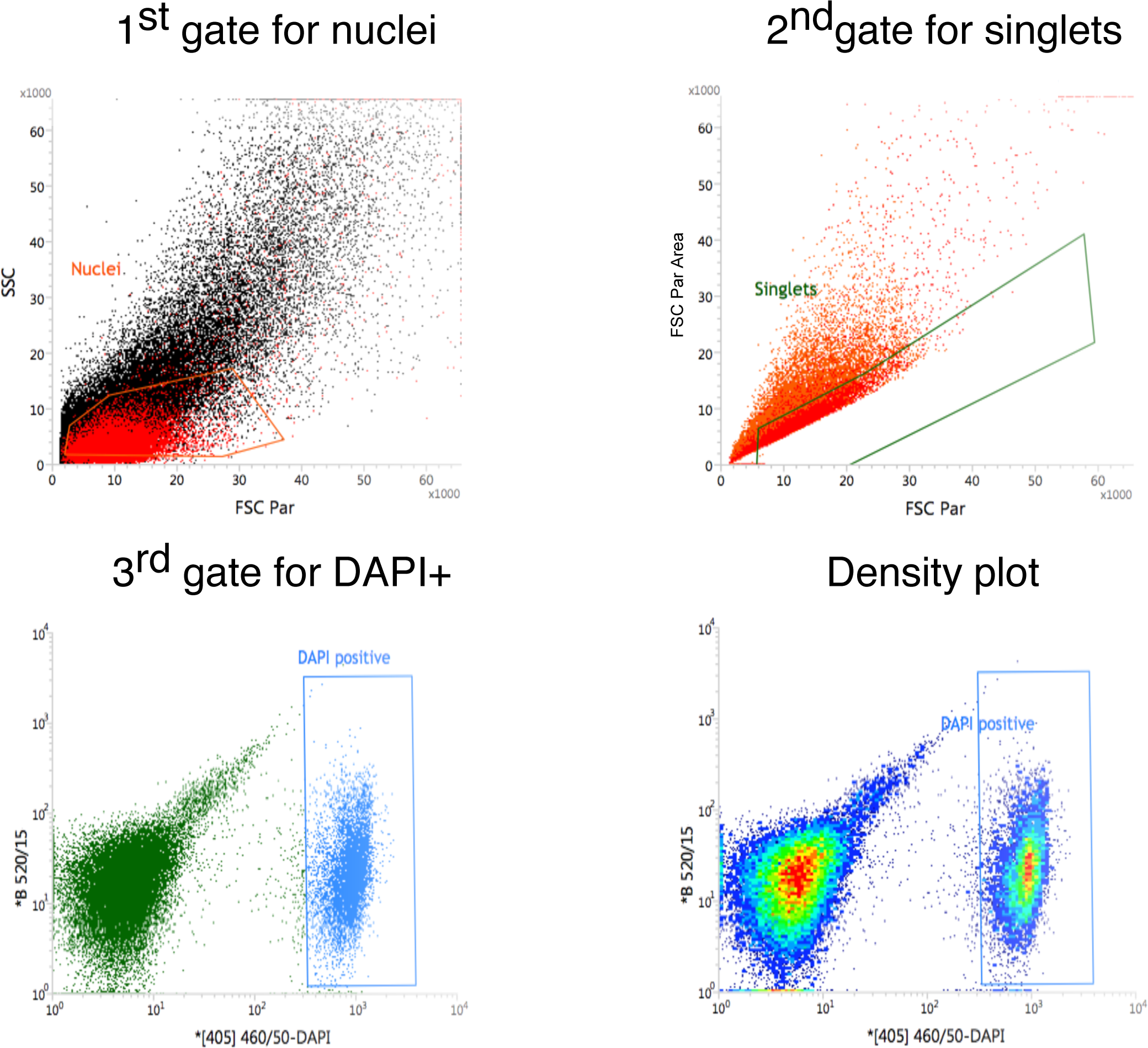
FANS gating strategy.

**Supplementary Fig. 12.**
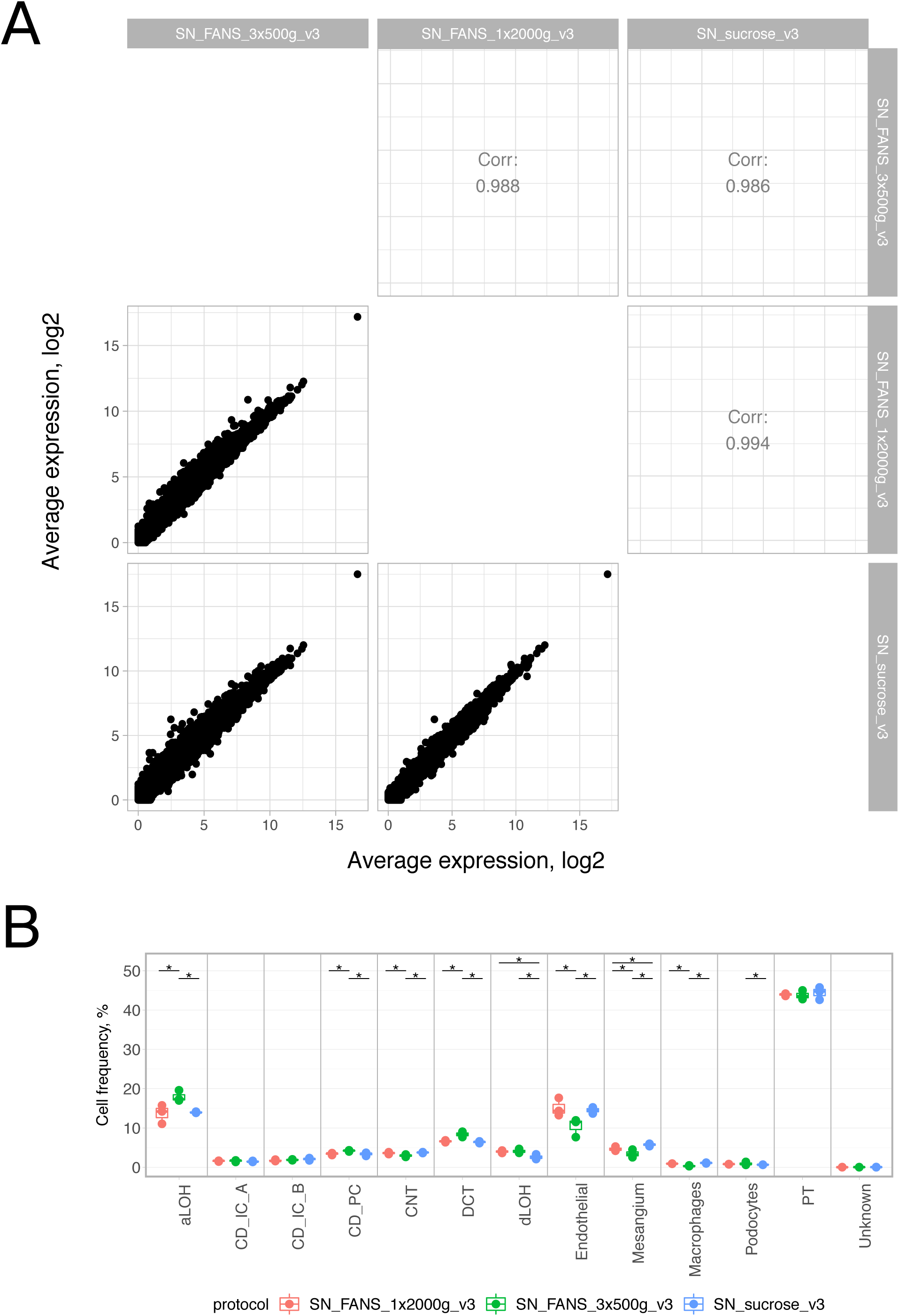
Comparison of nuclei isolation protocols. **a**) Aggregate gene expression and correlation between protocols. Raw counts were summed up for nuclei in each sample separately, then normalised to reads per million, averaged across biological replicates and log2-transformed with a pseudo count of 1 for plotting. Spearman correlation coefficients are shown. **b**) Cell type composition of the three nuclei isolation protocols. Three biological replicates are shown per protocol. Asterisks denote two-sided chi-square test p-value < 0.001.

**Supplementary Fig. 13.**
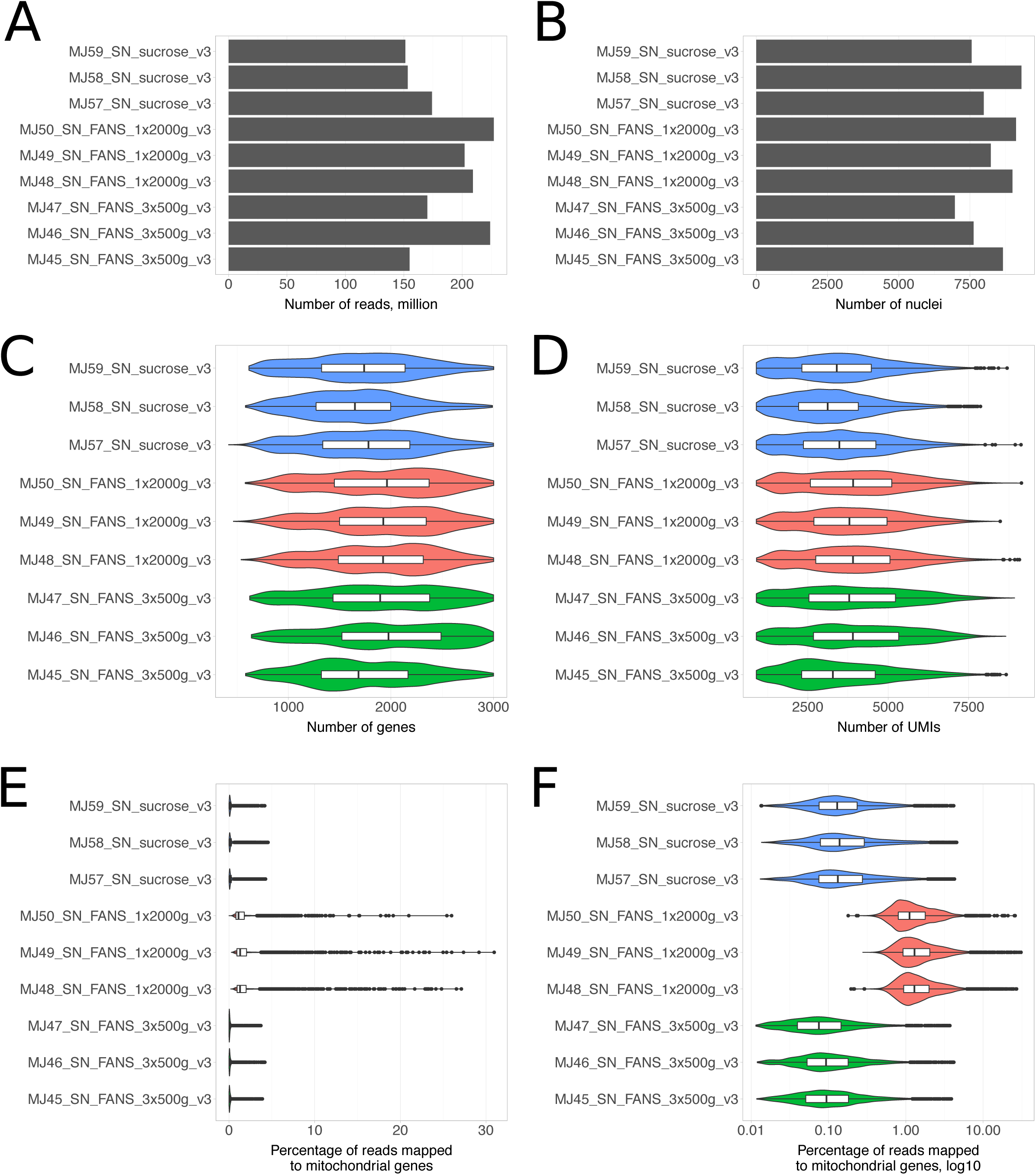
Comparison of nuclei isolation protocols. **a**) Total number of sequenced reads for each library. **b**) Number of nuclei passing all filtering steps as described in **Methods**. **c**) Distribution of the number of genes detected per nuclei. **d**) Distribution of the number of unique molecular identifiers detected per nuclei. **e**), **f**) Distribution of the percentage of reads mapped to mitochondrial genes per nuclei.

**Supplementary Fig. 14.**
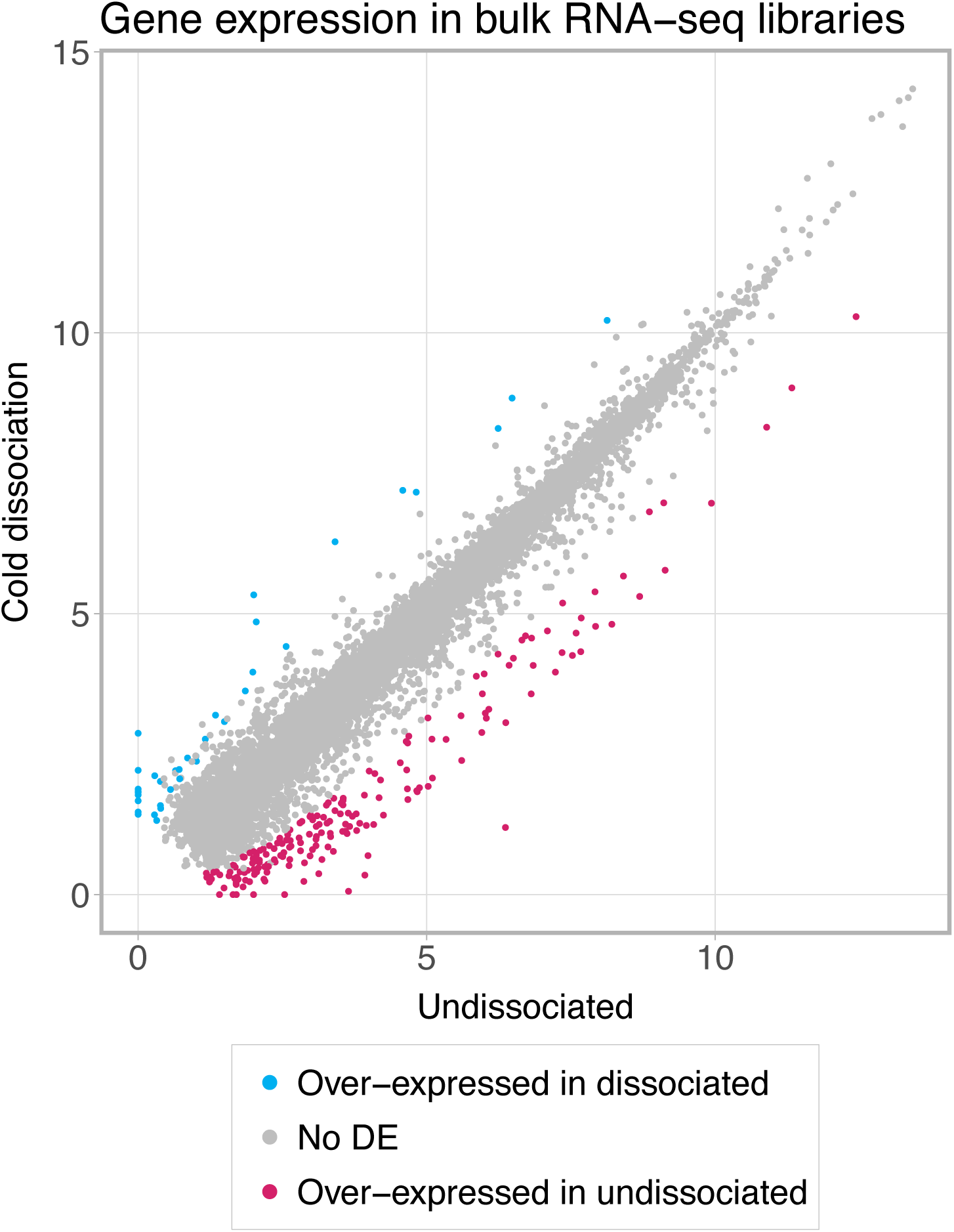
Comparison of bulk RNA-seq profiles of intact kidneys and cold-dissociated single-cell suspensions. GeTMM-normalised counts [43] were averaged across three biological replicates and log2-transformed after adding a pseudo count of 1. DEGs identified with FDR < 0.05 and logFC threshold of 2 using edgeR exact test [27] are indicated.

**Supplementary Fig. 15.**
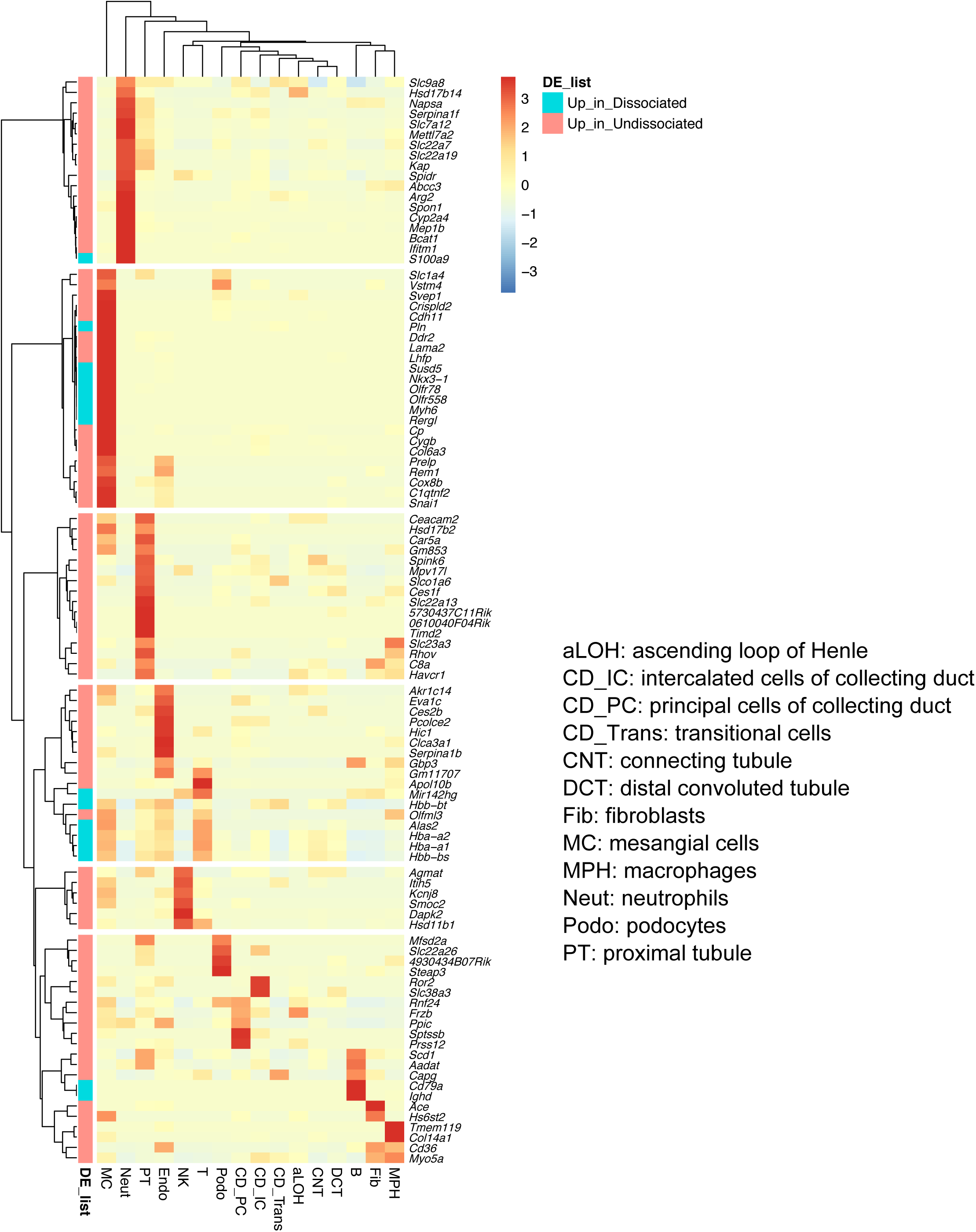
Expression of genes differentially expressed between bulk RNA-seq profiles of intact and dissociated kidneys in the matching single-cell dataset, Balb/c female mice. Normalised counts were averaged for each cell type, rows were scaled for plotting.

Supplementary Note 1. Microfluidics may alter cell composition.

To determine whether microfluidic partitioning could affect cell composition, we compared bulk RNA-seq and scRNA-seq data generated on the same dissociated kidney samples (**Fig. 1A, B**). We used BSEQ-sc [1] to predict the proportions of each cell type present in the samples before they were loaded on the microfluidics. Most notably, BSEQ-sc predicted cells of the ascending loop of Henle (aLOH) to be present at 18.6% and 14.3% on average for 3 biological replicates for warm and cold dissociation, respectively (**Supplementary Fig. 4**), while they were only present at 4.99% and 2.52% in the scRNA-seq libraries (warm- and cold-dissociated, respectively). Given that cells of aLOH are thought to be the second most populous cell type in kidney (estimated at 23.71% of kidney epithelial cells [2]), the analysis suggests the BSEQ-sc estimates are more likely to be correct and that aLOH cells may be lost in the microfluidic partitioning.

Podocytes were highly depleted in the warm dissociated samples (0.03% in warm vs 2.78% in cold); however, BSEQ-sc predicted podocytes to be at more similar abundance in the warm- and cold-dissociated samples (0.86% and 1.36%, respectively). In addition, known podocyte markers *Nphs1* and *Nphs2* were not significantly differentially expressed between bulk RNA-seq profiles of warm- and cold-dissociated kidneys (logFC = 0.397, FDR = 0.039 and logFC = 0.390, FDR = 0.032, respectively, edgeR exact test [3]). Together, this suggests that microfluidic partitioning likely contributes to the depletion of podocytes specifically in warm-dissociated kidneys.

Supplementary Note 2. Comparison of three nuclei isolation protocols.

Aggregate gene expression was highly correlated in the three nuclei isolation protocols (**Supplementary Fig. 12**) and each yielded similar numbers of nuclei with similar numbers of genes and UMIs detected per nucleus (**Supplementary Fig. 13**). The major difference observed was a higher percentage of reads mapping to mitochondrial genes in the SN_FANS_1x2000g_v3 data (mean of 1.69%), vs 0.27% and 0.15% for the SN_sucrose and SN_FANS_3x500g_v3 data, respectively (**Supplementary Fig. 13**). **Supplementary Tables 14-16** list differentially expressed genes for three pair-wise comparisons between the protocols, calculated for each cell type separately using Wilcoxon test in Seurat [4] with thresholds of logFC = 0.5, minimum detection rate 0.5, FDR < 0.05. The differential expression analysis suggested that contamination of all cell populations with highly expressed kidney transcripts was the strongest in SN_FANS_1x2000g_v3 and the lowest in SN_FANS_3x500g_v3. In terms of cell type composition, SN_FANS_1x2000g_v3 and SN_sucrose were mostly similar, while SN_FANS_3x500g showed significant differences across several cell populations (**Supplementary Fig. 12**). Taking the low levels of contamination and the simpler protocol without need for specialist FANS equipment we recommend the SN_sucrose protocol.

Supplementary Note 3. Comparison of bulk RNA-seq of intact kidneys and cold-dissociated cell suspensions.

We compared bulk RNA-seq profiles of undissociated kidneys to bulk RNA-seq profiles of cold-dissociated single-cell suspensions derived from kidneys of Balb/c female mice. Raw counts were normalised to gene length and then to library sizes using weighted trimmed mean of M-values (TMM) method in edgeR [3], to derive gene length corrected trimmed mean of M-values (GeTMM) as described in [5] (see **Methods**). Note that single-cell suspensions were filtered through 70µm and 40µm cell strainers, hence, this comparison could potentially reveal poorly dissociated cell types.

Differential expression analysis (edgeR exact test [3] with FDR < 0.05 and logFC threshold of 2) identified 191 genes with higher expression in undissociated kidneys and 36 genes with higher expression in dissociated kidneys (**Supplementary Fig. 14**, **Supplementary Table 18**). To get insight into the potential source of these genes, we investigated their expression levels in our single-cell dataset derived from the same batch of mice. Of a total of 227 differentially expressed genes, 26 genes were absent from the 10x transcriptome reference, hence, were not measured in single-cell experiments. Another 99 genes were not detected in the single-cell dataset, which could indicate either loss of certain cell types or low expression levels of these genes below the detection limit on a single-cell level. The latter is supported by the fact that genes which were detected in single-cell experiments had higher expression levels in undissociated bulk RNA-seq samples than genes not detected in single-cell experiments (mean expression of 200.9 versus 9.3 GeTMM-normalised counts, median 24.5 versus 4.1 GeTMM-normalised counts; two-sided Mann-Whitney test W = 7687, p-value < 2.2e-16). On the other hand, amongst the 99 genes not detected in the single-cell data, several are indicative of specific cell types. For example, the nervous tissue transcripts *Cck, Ak5* and *Gabra3* were more abundant in the intact kidney and may indicate RNA from nerve fibers which would not be expected to be seen in single-cell preparations.

Amongst the 102 differentially expressed genes which were detected in the single-cell dataset, 86 genes showed higher expression in undissociated kidneys and 16 – higher expression in dissociated suspensions. Amongst the 16 genes more abundant in the dissociated samples we identified haemoglobins (*Hbb-bs*, *Hba-a1*, *Hbb-bt*, *Hba-a2*) indicative of red blood cells. **Supplementary Fig. 15** shows a heatmap for average expression levels of the 102 genes in all cell types in the single-cell dataset. The heatmap reveals that several cell types, such as neutrophils, proximal tubules or endothelial cells, express genes which showed higher expression in undissociated than dissociated kidneys. This might indicate that the corresponding cell types were incompletely dissociated. Surprisingly, mesangial cells express both genes which showed higher expression in undissociated kidneys and genes with higher expression in dissociated suspensions, which might point to mesangial cell subtypes that are unequally represented in intact vs dissociated bulk RNA-seq kidney profiles.

## References

1. Cao J, Spielmann M, Qiu X, Huang X, Ibrahim DM, Hill AJ, et al. The single-cell transcriptional landscape of mammalian organogenesis. Nature. 2019;566:496–502.

2. Hochane M, van den Berg PR, Fan X, Berenger-Currias N, Adegeest E, Bialecka M, et al. Single-cell transcriptomics reveals gene expression dynamics of human fetal kidney development. PLoS Biol. 2019;17:e3000152.

3. Combes AN, Phipson B, Lawlor KT, Dorison A, Patrick R, Zappia L, et al. Single cell analysis of the developing mouse kidney provides deeper insight into marker gene expression and ligand-receptor crosstalk. Development. 2019;146.

4. Aizarani N, Saviano A, Sagar, Mailly L, Durand S, Herman JS, et al. A human liver cell atlas reveals heterogeneity and epithelial progenitors. Nature. 2019;572:199–204.

5. Single-cell transcriptomics of 20 mouse organs creates a Tabula Muris. Nature. 2018;562:367–72.

6. Lukowski SW, Lo CY, Sharov AA, Nguyen Q, Fang L, Hung SS, et al. A single-cell transcriptome atlas of the adult human retina. Embo j. 2019;38:e100811.

7. Puram SV, Tirosh I, Parikh AS, Patel AP, Yizhak K, Gillespie S, et al. Single-Cell Transcriptomic Analysis of Primary and Metastatic Tumor Ecosystems in Head and Neck Cancer. Cell. 2017;171:1611–24.e24.

8. Tirosh I, Izar B, Prakadan SM, Wadsworth MH, 2nd, Treacy D, Trombetta JJ, et al. Dissecting the multicellular ecosystem of metastatic melanoma by single-cell RNA-seq. Science. 2016;352:189–96.

9. Kim KT, Lee HW, Lee HO, Song HJ, Jeong da E, Shin S, et al. Application of single-cell RNA sequencing in optimizing a combinatorial therapeutic strategy in metastatic renal cell carcinoma. Genome Biol. 2016;17:80.

10. Li H, Courtois ET, Sengupta D, Tan Y, Chen KH, Goh JJL, et al. Reference component analysis of single-cell transcriptomes elucidates cellular heterogeneity in human colorectal tumors. Nat Genet. 2017;49:708–18.

11. Zanini F, Pu SY, Bekerman E, Einav S, Quake SR. Single-cell transcriptional dynamics of flavivirus infection. Elife. 2018;7.

12. Martin JC, Chang C, Boschetti G, Ungaro R, Giri M, Grout JA, et al. Single-Cell Analysis of Crohn’s Disease Lesions Identifies a Pathogenic Cellular Module Associated with Resistance to Anti-TNF Therapy. Cell. 2019;178:1493–508.e20.

13. Klein AM, Mazutis L, Akartuna I, Tallapragada N, Veres A, Li V, et al. Droplet barcoding for single-cell transcriptomics applied to embryonic stem cells. Cell. 2015;161:1187–201.

14. Macosko EZ, Basu A, Satija R, Nemesh J, Shekhar K, Goldman M, et al. Highly Parallel Genome-wide Expression Profiling of Individual Cells Using Nanoliter Droplets. Cell. 2015;161:1202–14.

15. Zheng GX, Terry JM, Belgrader P, Ryvkin P, Bent ZW, Wilson R, et al. Massively parallel digital transcriptional profiling of single cells. Nat Commun. 2017;8:14049.

16. van den Brink SC, Sage F, Vertesy A, Spanjaard B, Peterson-Maduro J, Baron CS, et al. Single-cell sequencing reveals dissociation-induced gene expression in tissue subpopulations. Nat Methods. 2017;14:935–6.

17. Potter SS. Single-cell RNA sequencing for the study of development, physiology and disease. Nat Rev Nephrol. 2018;14:479–92.

18. Adam M, Potter AS, Potter SS. Psychrophilic proteases dramatically reduce single-cell RNA-seq artifacts: a molecular atlas of kidney development. Development. 2017;144:3625–32.

19. Lake BB, Ai R, Kaeser GE, Salathia NS, Yung YC, Liu R, et al. Neuronal subtypes and diversity revealed by single-nucleus RNA sequencing of the human brain. Science. 2016;352:1586–90.

20. Krishnaswami SR, Grindberg RV, Novotny M, Venepally P, Lacar B, Bhutani K, et al. Using single nuclei for RNA-seq to capture the transcriptome of postmortem neurons. Nat Protoc. 2016;11:499–524.

21. Alles J, Karaiskos N, Praktiknjo SD, Grosswendt S, Wahle P, Ruffault PL, et al. Cell fixation and preservation for droplet-based single-cell transcriptomics. BMC Biol. 2017;15:44.

22. Wohnhaas CT, Leparc GG, Fernandez-Albert F, Kind D, Gantner F, Viollet C, et al. DMSO cryopreservation is the method of choice to preserve cells for droplet-based single-cell RNA sequencing. Sci Rep. 2019;9:10699.

23. Guillaumet-Adkins A, Rodriguez-Esteban G, Mereu E, Mendez-Lago M, Jaitin DA, Villanueva A, et al. Single-cell transcriptome conservation in cryopreserved cells and tissues. Genome Biol. 2017;18:45.

24. Bakken TE, Hodge RD, Miller JA, Yao Z, Nguyen TN, Aevermann B, et al. Single-nucleus and single-cell transcriptomes compared in matched cortical cell types. PLoS One. 2018;13:e0209648.

25. Lake BB, Codeluppi S, Yung YC, Gao D, Chun J, Kharchenko PV, et al. A comparative strategy for single-nucleus and single-cell transcriptomes confirms accuracy in predicted cell-type expression from nuclear RNA. Sci Rep. 2017;7:6031.

26. Wu H, Kirita Y, Donnelly EL, Humphreys BD. Advantages of Single-Nucleus over Single-Cell RNA Sequencing of Adult Kidney: Rare Cell Types and Novel Cell States Revealed in Fibrosis. J Am Soc Nephrol. 2019;30:23–32.

27. Robinson MD, McCarthy DJ, Smyth GK. edgeR: a Bioconductor package for differential expression analysis of digital gene expression data. Bioinformatics. 2010;26:139–40.

28. Chen J, Bardes EE, Aronow BJ, Jegga AG. ToppGene Suite for gene list enrichment analysis and candidate gene prioritization. Nucleic Acids Res. 2009;37:W305–11.

29. Hou R, Denisenko E, Forrest ARR. scMatch: a single-cell gene expression profile annotation tool using reference datasets. Bioinformatics. 2019.

30. Park J, Shrestha R, Qiu C, Kondo A, Huang S, Werth M, et al. Single-cell transcriptomics of the mouse kidney reveals potential cellular targets of kidney disease. Science. 2018;360:758–63.

31. Karaiskos N, Rahmatollahi M, Boltengagen A, Liu H, Hoehne M, Rinschen M, et al. A Single-Cell Transcriptome Atlas of the Mouse Glomerulus. J Am Soc Nephrol. 2018;29:2060–8.

32. Clark JZ, Chen L, Chou CL, Jung HJ, Lee JW, Knepper MA. Representation and relative abundance of cell-type selective markers in whole-kidney RNA-Seq data. Kidney Int. 2019;95:787–96.

33. Baron M, Veres A, Wolock SL, Faust AL, Gaujoux R, Vetere A, et al. A Single-Cell Transcriptomic Map of the Human and Mouse Pancreas Reveals Inter- and Intra-cell Population Structure. Cell Syst. 2016;3:346–60.e4.

34. Butler A, Hoffman P, Smibert P, Papalexi E, Satija R. Integrating single-cell transcriptomic data across different conditions, technologies, and species. Nat Biotechnol. 2018;36:411–20.

35. O’Sullivan ED, Mylonas KJ, Hughes J, Ferenbach DA. Complementary Roles for Single-Nucleus and Single-Cell RNA Sequencing in Kidney Disease Research. J Am Soc Nephrol. 2019;30:712–3.

36. Slyper M, Porter CBM, Ashenberg O, Waldman J, Drokhlyansky E, Wakiro I, et al. A single-cell and single-nucleus RNA-seq toolbox for fresh and frozen human tumors. bioRxiv. 2019:761429.

37. Chen J, Cheung F, Shi R, Zhou H, Lu W. PBMC fixation and processing for Chromium single-cell RNA sequencing. J Transl Med. 2018;16:198.

38. Martelotto L. ’Frankenstein’ protocol for nuclei isolation from fresh and frozen tissue for snRNA-seq. Protocolsio. 2019.

39. 10xGenomics; Isolation of nuclei for single cell RNA sequencing. 2017.

40. Lassmann T. TagDust2: a generic method to extract reads from sequencing data. BMC Bioinformatics. 2015;16:24.

41. Dobin A, Davis CA, Schlesinger F, Drenkow J, Zaleski C, Jha S, et al. STAR: ultrafast universal RNA-seq aligner. Bioinformatics. 2013;29:15–21.

42. Liao Y, Smyth GK, Shi W. featureCounts: an efficient general purpose program for assigning sequence reads to genomic features. Bioinformatics. 2014;30:923–30.

43. Smid M, Coebergh van den Braak RRJ, van de Werken HJG, van Riet J, van Galen A, de Weerd V, et al. Gene length corrected trimmed mean of M-values (GeTMM) processing of RNA-seq data performs similarly in intersample analyses while improving intrasample comparisons. BMC Bioinformatics. 2018;19:236.

44. Lun ATL, Riesenfeld S, Andrews T, Dao TP, Gomes T, Marioni JC. EmptyDrops: distinguishing cells from empty droplets in droplet-based single-cell RNA sequencing data. Genome Biol. 2019;20:63.

45. McCarthy DJ, Campbell KR, Lun AT, Wills QF. Scater: pre-processing, quality control, normalization and visualization of single-cell RNA-seq data in R. Bioinformatics. 2017;33:1179–86.

46. North BV, Curtis D, Sham PC. A note on the calculation of empirical P values from Monte Carlo procedures. Am J Hum Genet. 2002;71:439–41.

47. Benjamini Y, Hochberg Y. Controlling the False Discovery Rate: A Practical and Powerful Approach to Multiple Testing. Journal of the Royal Statistical Society: Series B (Methodological). 1995;57:289–300.

48. Rohrwasser A, Ishigami T, Gociman B, Lantelme P, Morgan T, Cheng T, et al. Renin and kallikrein in connecting tubule of mouse. Kidney Int. 2003;64:2155–62.

## References

1. Baron M, Veres A, Wolock SL, Faust AL, Gaujoux R, Vetere A, et al. A Single-Cell Transcriptomic Map of the Human and Mouse Pancreas Reveals Inter- and Intra-cell Population Structure. Cell Syst. 2016;3:346–60.e4.

2. Clark JZ, Chen L, Chou CL, Jung HJ, Lee JW, Knepper MA. Representation and relative abundance of cell-type selective markers in whole-kidney RNA-Seq data. Kidney Int. 2019;95:787–96.

3. Robinson MD, McCarthy DJ, Smyth GK. edgeR: a Bioconductor package for differential expression analysis of digital gene expression data. Bioinformatics. 2010;26:139–40.

4. Butler A, Hoffman P, Smibert P, Papalexi E, Satija R. Integrating single-cell transcriptomic data across different conditions, technologies, and species. Nat Biotechnol. 2018;36:411–20.

5. Smid M, Coebergh van den Braak RRJ, van de Werken HJG, van Riet J, van Galen A, de Weerd V, et al. Gene length corrected trimmed mean of M-values (GeTMM) processing of RNA-seq data performs similarly in intersample analyses while improving intrasample comparisons. BMC Bioinformatics. 2018;19:236.

